# Molecular Evolution across Mouse Spermatogenesis

**DOI:** 10.1101/2021.08.04.455131

**Authors:** Emily E. K. Kopania, Erica L. Larson, Colin Callahan, Sara Keeble, Jeffrey M. Good

## Abstract

Genes involved in spermatogenesis tend to evolve rapidly, but we still lack a clear understanding of how different components of molecular evolution vary across this complex developmental process. We used fluorescence activated cell sorting (FACS) to generate expression data for both early (meiotic) and late (postmeiotic) cell types across thirteen inbred strains of mice (*Mus*) spanning ~7 million years of evolution. We used these comparative developmental data to investigate the evolution of lineage-specific expression, protein-coding sequences, and expression levels. We found increased lineage specificity and more rapid protein-coding and expression divergence during late spermatogenesis, suggesting that signatures of rapid testis molecular evolution are punctuated across sperm development. Despite strong overall developmental parallels in these components of molecular evolution, protein and expression divergences were only weakly correlated across genes. We detected more rapid protein evolution on the X chromosome relative to the autosomes, while X-linked gene expression tended to be relatively more conserved likely reflecting chromosome-wide regulatory constraints. Using allele-specific FACS expression data from crosses between four strains, we found that the relative contributions of different regulatory mechanisms also differed between cell-types. Genes showing *cis*-regulatory changes were more common late in spermatogenesis, and tended to be associated with larger differences in expression levels and greater expression divergence between species. In contrast, genes with *trans*-acting changes were more common early and tended to be more conserved across species. Our findings advance understanding of gene evolution across spermatogenesis and underscore the fundamental importance of developmental context in molecular evolutionary studies.

## Introduction

Mature sperm are the most morphologically diverse animal cell type, likely as a consequence of intense selection on sperm form and function (Pitnick, et al. 2009). Genes involved in spermatogenesis also tend to evolve rapidly (Swanson, et al. 2003; Good and Nachman 2005; Turner, et al. 2008; Larson, et al. 2016; Finseth and Harrison 2018), suggesting that pervasive sexual selection also shapes molecular evolution (Swanson and Vacquier 2002; Harrison, et al. 2015). However, direct genotype-to-phenotype connections remain elusive for primary sexually selected traits, and there are additional evolutionary forces acting during spermatogenesis that shape overall patterns of molecular evolution (Good and Nachman 2005; Burgoyne, et al. 2009; Dean, et al. 2009; Larson, et al. 2016; Schumacher and Herlyn 2018). For example, many spermatogenesis genes are highly specialized (Eddy 2002; Chalmel, et al. 2007; Green, et al. 2018), which can relax pleiotropic constraint and contribute to rapid evolution even in the absence of positive directional selection (Winter, et al. 2004; Larracuente, et al. 2008; Meisel 2011). Other components of spermatogenesis are highly conserved because small disruptions can lead to infertility (Burgoyne, et al. 2009). Thus, spermatogenesis genes are likely to experience strong and sometimes contradictory evolutionary pressures. Understanding how these processes interact to shape molecular evolution across spermatogenesis is essential to understanding how natural selection shapes the genetic determinants of male fertility.

There are many components or levels of molecular evolution, spanning from protein sequence changes to differences in gene expression level, timing, and developmental specificity (King and Wilson 1975; Wray, et al. 2003; Larracuente, et al. 2008; Kaessmann 2010; Piasecka, et al. 2013; Cridland, et al. 2020). Many of these components have been shown to evolve relatively rapidly during spermatogenesis (Meiklejohn, et al. 2003; Khaitovich, et al. 2005; Voolstra, et al. 2007; Brawand, et al. 2011; Harrison, et al. 2015; Vicens, et al. 2017; Cridland, et al. 2020; Sánchez-Ramírez, et al. 2021), and generally trend towards increased divergence during the later stages of development (Good and Nachman 2005; Piasecka, et al. 2013; Larson, et al. 2016). Novel genes disproportionately arise with testis-specific expression (Levine, et al. 2006; Zhao, et al. 2014; Schroeder, et al. 2019; Cridland, et al. 2020; Lange, et al. 2021), likely as a consequence of the more permissive regulatory environment of the later stages of sperm development (Kaessmann 2010; Soumillon, et al. 2013). Likewise, the later stages of spermatogenesis tend to be enriched for novel testis-specific genes (Eddy 2002; Chalmel, et al. 2007; Green, et al. 2018). These developmental signatures of novelty and specialization are further reflected in patterns of increased divergence of protein sequences (Good and Nachman 2005; Kousathanas, et al. 2014) and expression levels (Larson, et al. 2016) between species during the later stages of sperm development. Parallel signatures of rapid molecular evolution likely reflect both relaxed constraints during the late stages of spermatogenesis, and enhanced positive selection on late-developing sperm phenotypes (Eddy 2002; Good and Nachman 2005; Larracuente, et al. 2008; Larson, et al. 2016; Cutter and Bundus 2020). However, it remains unclear how strongly different forms of molecular evolution are correlated. For example, changes in gene expression may often be cell or stage-specific and therefore may be less pleiotropic than protein-coding changes. This pleiotropic constraint hypothesis primarily applies to *cis*-regulatory changes, which likely affect one gene, whereas *trans*-regulatory changes can affect many genes across multiple cell types (Wray, et al. 2003; Carroll 2008; Cutter and Bundus 2020).

The X chromosome provides a compelling example of how the conflicting selective pressures acting on spermatogenesis may shape different components of molecular evolution. Theory predicts that the X chromosome should evolve more rapidly than the autosomes due to a lower effective population size (i.e., increased genetic drift) and more efficient selection resulting from hemizygosity in males, which immediately exposes recessive beneficial and deleterious mutations to selection (Charlesworth, et al. 1987; Vicoso and Charlesworth 2009). Consistent with these predictions, protein-coding evolution tends to be faster on the X chromosome compared to the autosomes in several taxa, and this effect is often strongest for genes with male-biased expression (Khaitovich, et al. 2005; Baines and Harr 2007; Baines, et al. 2008; Meisel and Connallon 2013; Parsch and Ellegren 2013; Larson, et al. 2016). Novel genes tend to arise more often on the X chromosome, and these are often expressed during spermatogenesis (Levine, et al. 2006; Kaessmann 2010). There is also some evidence for rapid expression evolution on the X chromosome in flies and mammals (Khaitovich, et al. 2005; Brawand, et al. 2011; Meisel, et al. 2012; Coolon, et al. 2015), but X-linked expression in mice appears conserved relative to autosomal genes expressed during the later stages of spermatogenesis (Larson, et al. 2016). Stage-specific differences in relative rates of expression evolution on the X chromosome may result from the unique regulatory pattern that the sex chromosomes undergo during mammalian spermatogenesis. In males, the X chromosome is inactivated early in meiosis (i.e., meiotic sex chromosome inactivation, MSCI; McKee and Handel 1993) and remains partially repressed during the postmeiotic haploid stages of sperm development (i.e., postmeiotic sex chromosome repression, PSCR; Namekawa, et al. 2006).

These stage-specific patterns highlight the importance of studying specific components of molecular evolution in a developmental framework (fig. 1A; Larson, et al. 2018a; Cutter and Bundus 2020). However, studies of molecular evolution have primarily focused on pairwise contrasts across nuanced aspects of tissue development (Good and Nachman 2005; Larson, et al. 2016), or examined protein-coding versus regulatory evolution in whole tissues (Khaitovich, et al. 2005; Voolstra, et al. 2007; Larson, et al. 2016; Mack, et al. 2016; Vicens, et al. 2017; Cridland, et al. 2020), without combining both in a phylogenetic framework. Relying on whole tissue expression comparisons may be particularly problematic for spermatogenesis, because differences in testis composition are expected to evolve rapidly between species (Ramm and Schärer 2014; Yapar, et al. 2021) and may confound patterns of expression level divergence (Good, et al. 2010; Larson, et al. 2016; Hunnicutt, et al. 2021). Nonetheless, collection of stage or cell-specific expression data remains technically demanding (da Cruz, et al. 2016; Green, et al. 2018), likely limiting widespread use in comparative studies. As a consequence, most evolutionary studies of gene expression have relied on whole tissue comparisons between closely-related species pairs, instead of using more powerful phylogenetic approaches (Rohlfs and Nielsen 2015; Dunn, et al. 2018).

**Figure 1.**
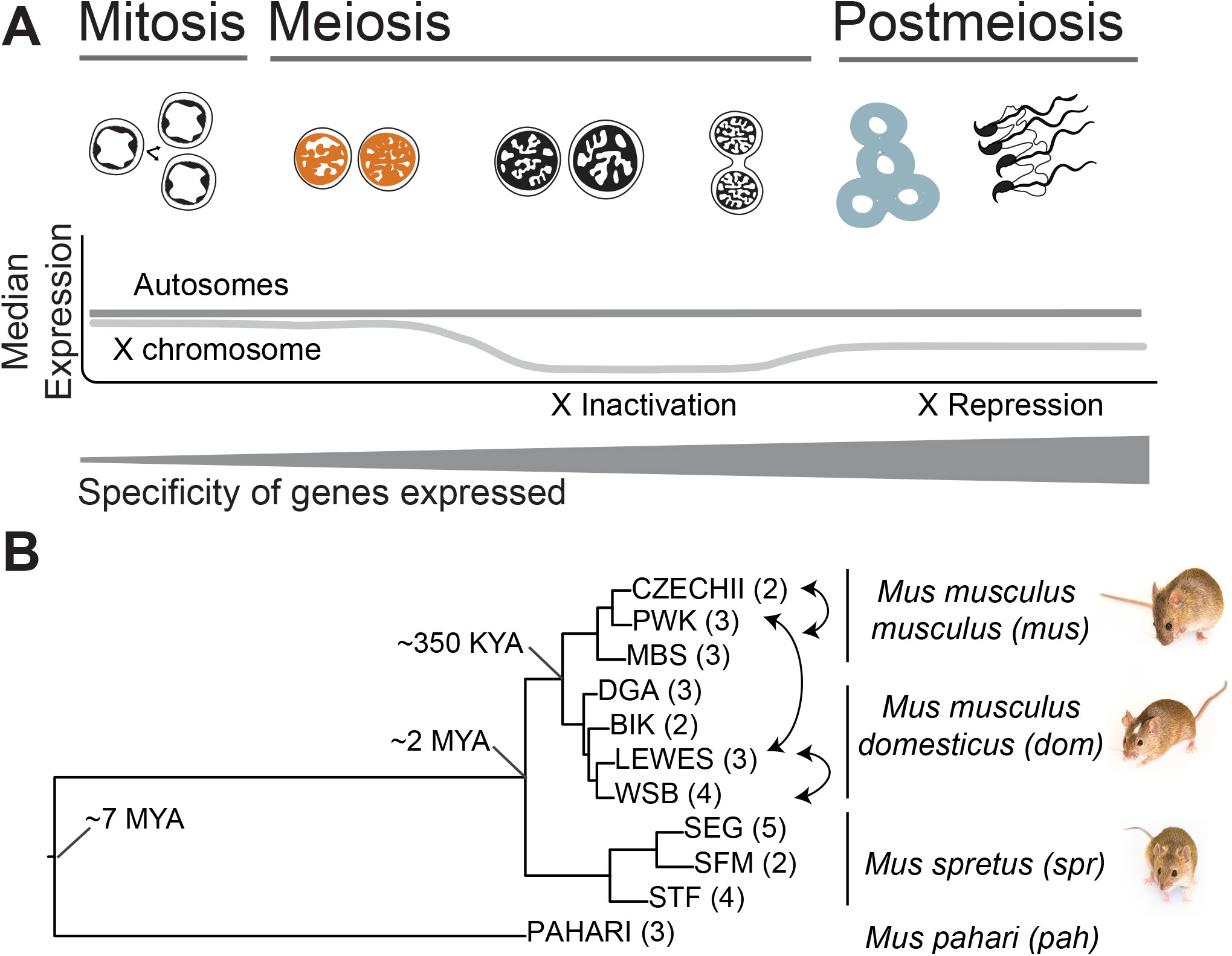
(*A*) Predictive framework depicting the major stages of spermatogenesis and expected relative expression levels of the X chromosome and autosomes at each stage (Namekawa, et al. 2006). The two cell populations used in this study are leptotene-zygotene (“early”, second from left, orange) and round spermatids (“late”, second from right, blue). The relative thickness of the gray bar represents the predicted cell type specificity at each stage (Eddy 2002; Chalmel, et al. 2007; Larson, et al. 2016; Green, et al. 2018). (*B*) Maximum likelihood tree of concatenated exome data from the four *Mus* species or subspecies used in this study: *Mus musculus musculus* (*mus*), *Mus musculus domesticus* (*dom*), *Mus spretus* (*spr*), *Mus pahari* (*pah*). Tips are labeled with the inbred strains from each lineage, with select crosses used to generate F1 hybrids indicated with arrows. Number of individuals sampled for each strain indicated in parentheses. Approximate divergence times are placed at each major node (Chevret, et al. 2005). All nodes had 100% bootstrap support.

In this study, we use a comparative developmental approach to gain a more comprehensive understanding of molecular evolution across spermatogenesis in house mice (*Mus*). Mice are the predominant laboratory model for mammalian reproduction (Phifer-Rixey and Nachman 2015; Firman 2020), with abundant genomic resources (Keane, et al. 2011; Thybert, et al. 2018), and established wild-derived inbred strains that can be crossed to resolve mechanisms underlying expression divergence (i.e., *cis*-versus *trans*-regulatory changes; Mack, et al. 2016). Mice also show divergence in sperm head morphologies across closely related species (Skinner, et al. 2019) and experience sperm competition in the wild (Dean, et al. 2006), providing a compelling system for understanding the evolution of spermatogenesis.

We used fluorescence activated cell sorting (FACS) to resolve patterns of gene expression in two enriched spermatogenic cell populations across several mouse strains, species, and cross-types (fig. 1A). Our study used two main comparisons. First, we evaluated divergence in spermatogenic protein sequences and gene expression levels across thirteen inbred strains of mice, including two subspecies of the house mouse (*Mus musculus*) and two other *Mus* species spanning seven million years of evolution (fig. 1B; Chevret, et al. 2005). Second, we used published data from reciprocal crosses between a subset of these inbred strains to resolve the relative contribution of *cis*-versus *trans*-regulatory changes to expression divergence. We used these data to address five main questions: (i) Is gene expression more lineage-specific during late spermatogenesis? (ii) Do protein-coding sequences and gene expression levels evolve faster during the later stages of spermatogenesis? (iii) Is the rate of molecular evolution elevated on the X chromosome compared to the autosomes, and does this relationship change across spermatogenesis? (iv) To what extent are protein-coding and gene expression divergence correlated, and does this relationship change across developmental stages? (v) Are there differences in the relative contributions of regulatory mechanisms (*cis*-versus *trans*-regulatory changes) across spermatogenesis?

## Results

### Spermatogenesis Gene Expression by Cell Type and Lineage

We collected spermatogenesis expression data from 34 mice representing four different species or subspecies: *Mus musculus musculus*, *Mus musculus domesticus*, *Mus spretus*, and *Mus pahari*. We will use the abbreviations *mus*, *dom*, *spr*, and *pah* to reference the four major groups, and refer to all taxa as “lineages” for concision (fig. 1B). For each sample, we generated expression data for two spermatogenic cell types, an early meiotic cell type (leptotene-zygotene cells from early prophase of meiosis I, hereafter “early”) and a post-meiotic cell type (round spermatids, hereafter “late”). We identified 23,164 one-to-one orthologs, including both protein-coding and non-protein-coding genes, that were annotated in all four mouse lineages and the mouse reference (GRCm38). From this set, we defined expressed genes as those with an FPKM > 1 in all samples of a given cell type. Expression variance cleanly separated samples by cell type and lineage (supplementary fig. S1, Supplementary Material online), indicating successful enrichment of different cell types. Most expressed genes were detected in both cell types (table 1). However, approximately one third of the detected genes were preferentially expressed or “induced” in a given cell type (transcripts with > 2X median expression level in one cell type across all lineages; table 1). We also identified expressed genes that show testis-specific expression based on published multi-tissue expression data (Chalmel, et al. 2007). We found that 493 testis-specific genes were induced late, while only 65 testis-specific genes were induced early (table 1), consistent with increased specificity late in spermatogenesis (Eddy 2002; Larson, et al. 2016; Green, et al. 2018). To distinguish experimental noise from biologically meaningful expression, we also used a Bayesian approach to determine if a gene was “active” in a tissue or cell type (Thompson, et al. 2020) and found broad overlap with genes in the expressed dataset (table 1). Using the same framework, we identified genes showing evidence for lineage-specific expression (“active” in a single lineage or subset of lineages).

**Table 1.** Counts of genes in each dataset and spermatogenic cell type. Numbers in parentheses represent the percent of genes in the “active” datasets that were also in the “expressed” dataset. TS = testis-specific (inferred from Chalmel, et al. 2007)

|  | early<br>(spermatocytes) | late<br>(round spermatids) | both early and late |
| --- | --- | --- | --- |
| expressed | 9570 | 8986 | 7670 |
| induced | 3375 | 2769 | 0 |
| testis-specific (TS) | 544 | 655 | 524 |
| induced and TS | 65 | 493 | 0 |
| active (*dom*) | 8206 (98.2%) | 8581 (90.4%) | 6355 |
| active (*mus*) | 8782 (97.5%) | 10098 (83.4%) | 7289 |
| active (*spr*) | 8728 (97.1%) | 9509 (86.0%) | 7227 |
| active (*pah*) | 8124 (97.6%) | 9563 (83.9%) | 6682 |

We found that lineage-specificity was rare overall, but more common for autosomal genes active during late spermatogenesis (Pearson’s χ2 test; *dom*: P << 0.0001, *mus*: P << 0.0001, *spr*: P << 0.0001, *dom*-*mus* common ancestor: P << 0.0001; fig. 2A). X-linked genes showed no significant differences in lineage-specificity between early and late cell types (fig. 2B), which could reflect a lack of specialization on the sex chromosomes, or reduced power to detect differences between cell types given small sample sizes. Few genes were lineage-specific in both cell types, and all were autosomal (*dom*: 9 genes, *mus*: 24 genes, *spr*: 24 genes, *dom*-*mus*: 21 genes). We found similar results using a log fold-change (logFC) approach to identify lineage-specific genes (supplementary fig. S2, Supplementary Material online). These results suggest that lineage-specific expression of spermatogenic genes is relatively uncommon at these shallow phylogenetic scales, but more likely to arise later in spermatogenesis.

**Figure 2.**
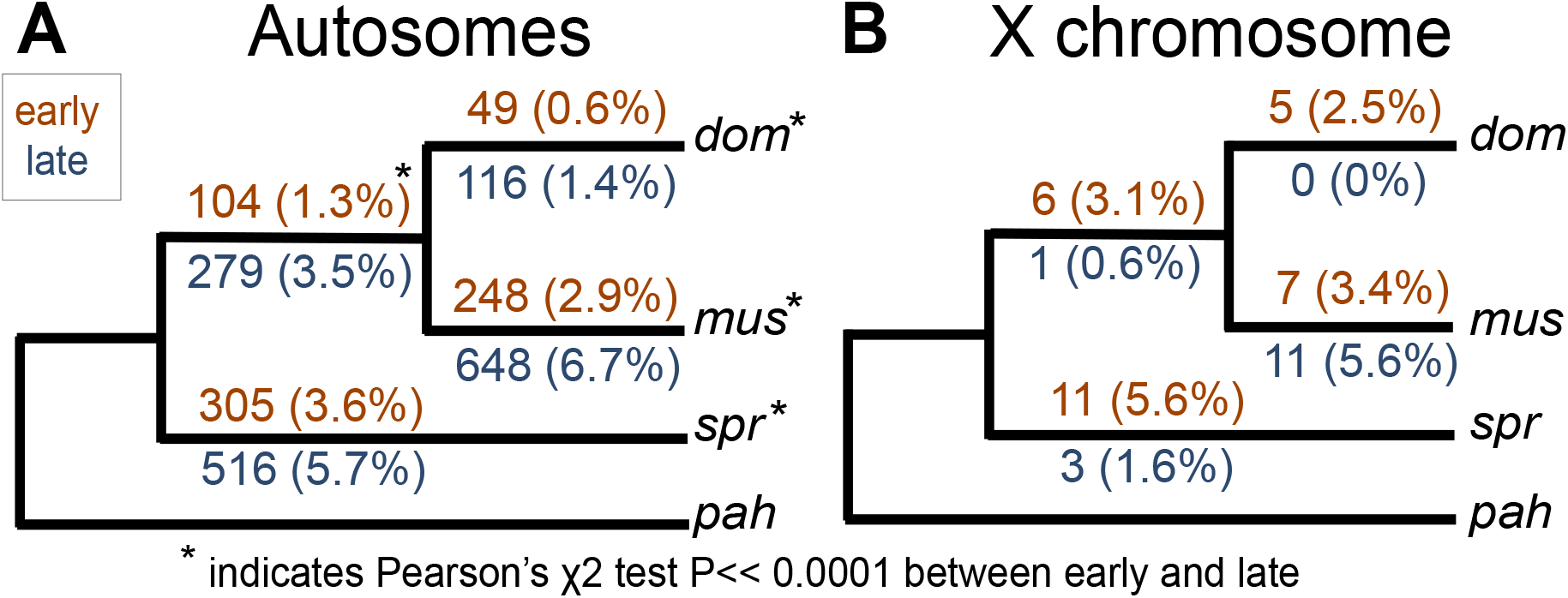
Number of genes that were lineage-specific on each internal branch of the mouse phylogeny used in this study. Numbers in parentheses are the percent of active genes that were lineage-specific. Results are presented separately for the autosomes (*A*) and X chromosome (*B*). Orange values above each branch represent the early cell type and blue values below represent the late cell type. Asterisks indicate a significant difference between early and late on that branch based on a Pearson’s χ2 test.

### Greater Protein-Coding and Gene Expression Divergence during Late Spermatogenesis

Having detected subtle increases in lineage-specificity late in spermatogenesis, we next tested if rates of protein sequence evolution (dN/dS) and expression level divergence were also elevated during the postmeiotic stage, as has been reported previously (Larson, et al. 2016). Genes induced late in spermatogenesis showed significantly higher rates of protein-coding divergence on both the autosomes (n = 2046 genes induced early, median dN/dS = 0.11; n = 1711 genes induced late, median dN/ dS = 0.20; Wilcoxon rank sum test P << 0.0001) and the X chromosome (n = 54 genes induced early, median dN/dS = 0.25; n = 61 genes induced late, median dN/dS = 0.41; Wilcoxon rank sum test P = 0.049; fig. 3A, supplementary tables S1 and S2, Supplementary Material online). The 489 testis-specific genes showed elevated dN/dS overall, but most testis-specific genes were expressed in both cell types and there was no significant difference between genes expressed early and late for the autosomes (n = 350 genes expressed early, median dN/dS = 0.28; n = 424 genes expressed late, median dN/dS = 0.30; Wilcoxon rank sum test P = 1) or the X chromosome (n = 16 genes expressed early; median dN/dS = 0.59; n = 24 genes expressed late, median dN/dS = 0.58; Wilcoxon rank sum test P = 1). However, 348 testis-specific genes were preferentially expressed in the late cell type, representing ~20% of all genes induced late for which we were able to calculate dN/dS. Taken together, these results confirm that tissue specificity plays an important role in the rapid protein-coding divergence of spermatogenic genes, and that most of this signature involves genes induced during postmeiotic spermatogenesis.

**Figure 3.**
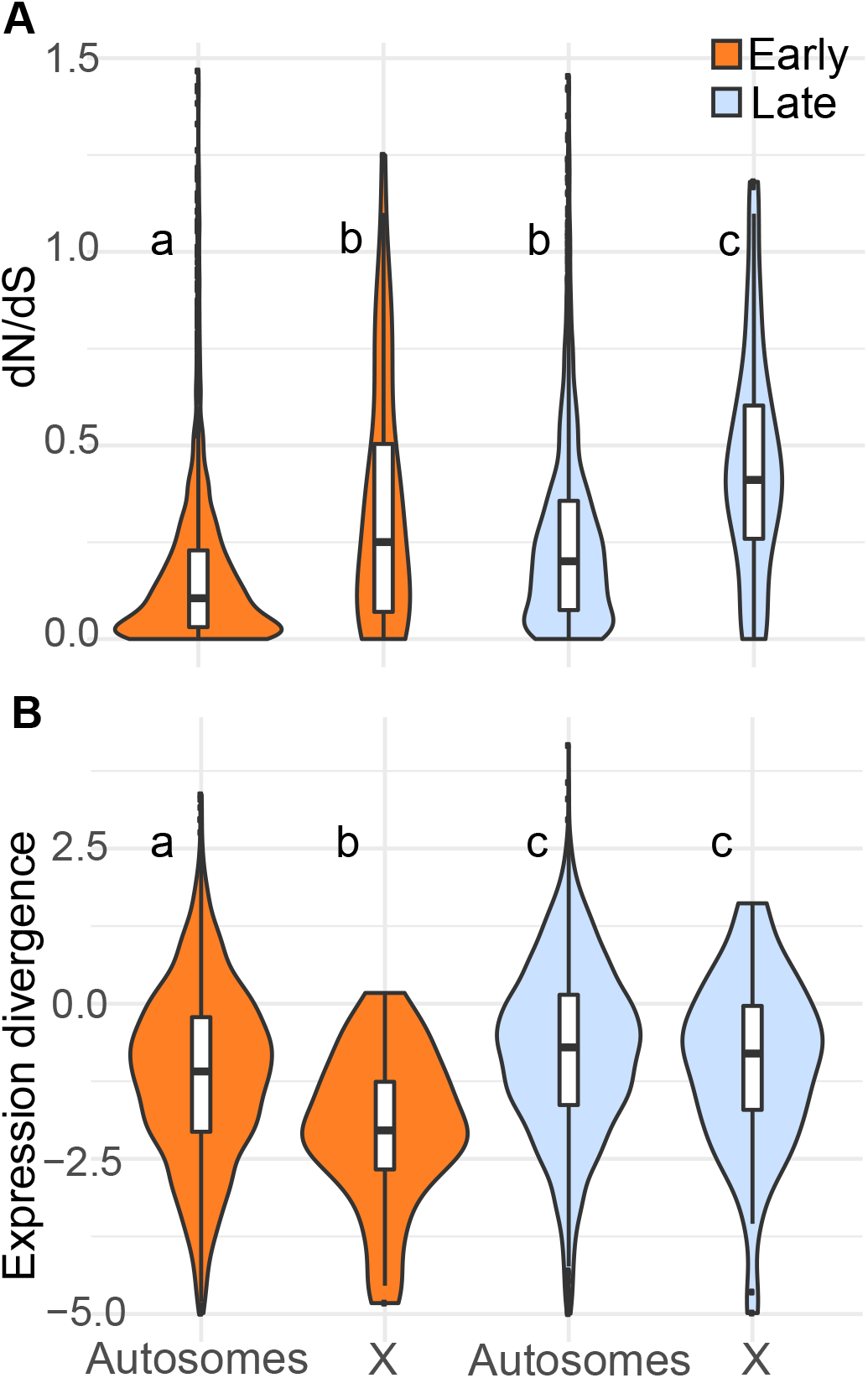
(*A*) Protein-coding and (*B*) expression divergence on the autosomes and X chromosome for genes induced in each cell type. Expression divergence values on the y-axis are −log(*beta*), where *beta* is the measure of expression divergence from EVE. Higher values on the y-axis represent higher divergence. The center of each violin plot is a standard boxplot, with the center horizontal line representing the median divergence value. The violins show the probability density of divergence values for each group. A wider part of the violin at a given value means genes expressed in that group are more likely to have that divergence value. The letters above each violin indicate significant differences between the cell types and chromosome types based on a Wilcoxon rank-sum test.

We used a phylogenetic ANOVA to estimate expression divergence while controlling for phylogenetic relatedness and variance within lineages [i.e., the Expression Variance and Evolution (EVE) model; Rohlfs and Nielsen 2015]. We report expression divergence from EVE as −log(*beta_i_*), where *beta_i_* is a metric from EVE that represents the ratio of within-lineage variance to between-lineage evolutionary divergence, and higher positive −log(*beta_i_*) values correspond to greater divergence between lineages.

Expression divergence was higher for genes induced late in spermatogenesis on both the autosomes (n = 2461 genes induced early, median EVE divergence = −1.09; n = 2305 genes induced late, median EVE divergence = −0.70; Wilcoxon rank sum test P << 0.0001) and X chromosome (n = 44 genes induced early, median EVE divergence = −2.04; n = 68 genes induced late, median EVE divergence = −0.80; Wilcoxon rank sum test P = 0.00019; fig. 3B). This pattern held for all expressed genes and testis-specific genes (supplementary table S3, Supplementary Material online). We also found higher divergence late for expressed and induced autosomal genes (supplementary table S3, Supplementary Material online) based on pairwise expression divergences using logFC and the metric from Meisel, et al. (2012); however, the pairwise framework did not give a consistent pattern on the X chromosome. When looking at all genes, most pairwise comparisons showed higher divergence late, but induced genes showed no difference between early and late spermatogenesis for most comparisons. However, the *dom* versus *spr* comparison had lower divergence late for all expressed genes and induced genes (supplementary table S3, Supplementary Material online). Collectively, we found strong evidence for more rapid protein-coding and gene expression level divergence during postmeiotic spermatogenesis, suggesting that these general patterns hold after controlling for phylogeny and at deeper divergence levels than had previously been shown in mice (Larson, et al. 2016). Despite our expanded phylogenetic sample, we still lacked the power to determine if more rapid expression and protein-coding divergence is due to positive directional selection (supplementary fig. S3, Supplementary Material online).

### Weak Positive Correlation between Gene Expression and Protein-Coding Divergence

We next tested for more general relationships between protein-coding and expression divergence across sets of genes expressed or induced during spermatogenesis (supplementary fig. S4, supplementary table S4, Supplementary Material online). Across all autosomal genes expressed early, there was a weak positive correlation between dN/dS and pairwise expression divergence (ρ = 0.13-0.17, Spearman’s rank correlation P << 0.0001). For induced genes, this correlation was weaker but still significant (ρ = 0.07-0.11, Spearman’s rank correlation P < 0.05). For the late cell type, there was also a weak positive correlation between pairwise expression divergence and dN/dS on the autosomes, but the correlation was weaker than that seen in the early cell type (ρ = 0.03-0.05, Spearman’s rank correlation P < 0.05). There was no correlation for the set of genes induced late. When looking only at genes with evidence for positive directional selection at the protein-coding level after correction for multiple tests (366 genes), the correlation was stronger on the autosomes late for the *dom* vs *spr* (n = 250 genes, ρ = 0.17, Spearman’s rank correlation P = 0.02) and *mus* vs *spr* comparisons (n = 249 genes, ρ = 0.18, Spearman’s rank correlation P << 0.0001). When comparing dN/dS to EVE expression divergence, we only saw a significant positive correlation for genes expressed late that were also under positive selection at the protein-coding level (n = 160 genes, ρ = 0.18, Spearman’s rank correlation P = 0.04). In summary, we tended to observe a positive relationship between protein-coding and expression level divergence, but the strength of this relationship was weak and varied by gene set and divergence metric.

### Faster-X Protein-Coding but Not Gene Expression Evolution

In addition to comparisons between spermatogenesis cell types, we compared relative rates of molecular evolution between X-linked and autosomal genes within a cell type. We found that protein-coding divergence was higher on the X chromosome, both early and late, across all gene sets (fig. 3A, supplementary tables S1 and S2, Supplementary Material online) consistent with several previous studies (Khaitovich, et al. 2005; Baines, et al. 2008; Meisel and Connallon 2013; Kousathanas, et al. 2014; Larson, et al. 2016). For expression evolution, we found lower divergence on the X chromosome early using EVE (n = 2461 autosomal genes, median EVE divergence = −1.09; n = 44 X-linked genes, median EVE divergence = −2.04; Wilcoxon rank sum test P =0.00015; fig. 3B), but higher X-linked divergence when using pairwise comparisons (supplementary table S3, Supplementary Material online). A major difference between these approaches was that EVE calculates divergence across a phylogeny, so genes that show divergent expression levels in one lineage may still be conserved across the entire phylogeny. We detected significant correlations between pairwise divergence values for different pairwise comparisons on the autosomes, and during late spermatogenesis, but lower or non-significant correlations on the X early (table 2). Thus, many genes on the X chromosome expressed early showed relatively high divergence between two particular lineages, but lower divergence across other pairwise comparisons and across the phylogeny as a whole. This lineage-specific variance underscores the importance of evaluating gene expression divergence in a phylogenetic framework (Rohlfs and Nielsen 2015; Dunn, et al. 2018).

**Table 2.** Correlation between pairwise expression divergence values for all possible pairwise comparisons. Numbers presented are ρ values from a Spearman’s rank correlation test. We tested for correlations in pairwise expression divergence value among induced genes in each stage and chromosome group (early X, early autosomal, late X, and late autosomal). Gray boxes indicate no significant correlation between pairwise divergence values after FDR correction (Spearman’s rank correlation P > 0.05). Bolded values indicate the lowest Spearman’s ρ value for each pairwise comparison across the four stages and chromosome groups.

|  |  | *dom* vs *mus* | *dom* vs *spr* | *mus* vs *spr* | *dom* vs *pah* | *mus* vs *pah* |
| --- | --- | --- | --- | --- | --- | --- |
| **Early, X-linked** | *dom* vs *spr* | 0.34 |  |  |  |  |
|  | *mus* vs *spr* | **0.07** | **0.28** |  |  |  |
|  | *dom* vs *pah* | **0.07** | **0.14** | **0.19** |  |  |
|  | *mus* vs *pah* | **0.16** | **0.10** | **0.03** | **0.62** |  |
|  | *spr* vs *pah* | **0.14** | 0.27 | **0.16** | 0.58 | 0.67 |
| **Early, autosomal** | *dom* vs *spr* | **0.32** |  |  |  |  |
|  | *mus* vs *spr* | 0.32 | 0.61 |  |  |  |
|  | *dom* vs *pah* | 0.28 | 0.28 | 0.27 |  |  |
|  | *mus* vs *pah* | 0.29 | 0.26 | 0.30 | 0.74 |  |
|  | *spr* vs *pah* | 0.24 | 0.32 | 0.34 | **0.55** | **0.57** |
| **Late, X-linked** | *dom* vs *spr* | 0.36 |  |  |  |  |
|  | *mus* vs *spr* | 0.50 | 0.45 |  |  |  |
|  | *dom* vs *pah* | 0.20 | 0.23 | 0.22 |  |  |
|  | *mus* vs *pah* | 0.28 | 0.28 | 0.36 | 0.74 |  |
|  | *spr* vs *pah* | 0.15 | **0.20** | 0.20 | 0.73 | 0.72 |
| **Late, autosomal** | *dom* vs *spr* | 0.35 |  |  |  |  |
|  | *mus* vs *spr* | 0.37 | 0.59 |  |  |  |
|  | *dom* vs *pah* | 0.30 | 0.33 | 0.30 |  |  |
|  | *mus* vs *pah* | 0.30 | 0.30 | 0.33 | 0.76 |  |
|  | *spr* vs *pah* | 0.25 | 0.32 | 0.33 | 0.64 | 0.63 |

In late spermatogenic cells (i.e., round spermatids), X-linked expression divergence was similar to or lower than on the autosomes depending on the contrast and approach. Using EVE, we found similar divergence on the X chromosome and autosomes late (n = 2305 autosomal genes, median EVE divergence = −0.70; n = 68 X-linked genes, median EVE divergence = −0.80; Wilcoxon rank sum test P = 0.34; fig. 3B), while pairwise comparisons gave mixed results, depending on which two lineages were compared (supplementary table S3, Supplementary Material online). There were proportionally fewer differentially expressed genes on the X chromosome (fig. 4, supplementary fig. S5, Supplementary Material online), and this pattern was strongest for the more closely related comparisons (hypergeometric test; *mus* versus *dom* P << 0.0001, *spr* versus *dom* P << 0.0001, *spr* versus *mus* P << 0.0001). Across all metrics of expression divergence and both developmental stages, there was no evidence for pervasive faster-X gene expression level evolution.

**Figure 4.**
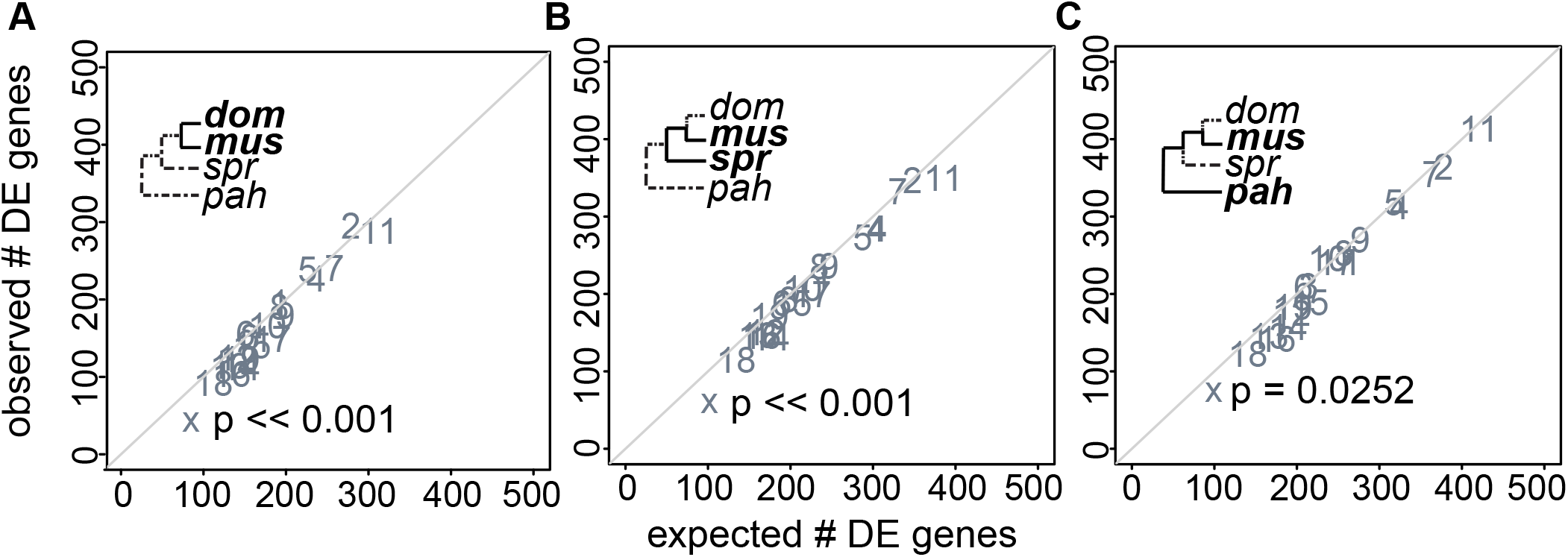
Observed versus expected number of genes differentially expressed (DE) in late spermatogenesis for three pairwise comparisons at different levels of evolutionary divergence: (*A*) *dom* versus *mus*, (*B*) *spr* versus *mus*, and (*C*) *pah* versus *mus*. Each point represents a different chromosome. The diagonal line is the one-to-one line at which the observed number of DE genes equals the expected number. P-values are shown for the X chromosome only. They are based on a hypergeometric test for enrichment and corrected for multiple tests using a false discovery rate correction. A significant p-value indicates that the observed number of DE genes is different from the expected number.

### Relative Contributions of cis- and trans-Regulatory Evolution Vary across Spermatogenesis

Having shown differences in expression divergence between cell types, we next asked if there were differences in the types of regulatory mutations (e.g., *cis*-versus *trans*-regulatory changes) underlying expression divergence in each cell type. We used whole testis (Mack, et al. 2016) and FACS-sorted (Larson, et al. 2017) data from reciprocal crosses between house mouse subspecies (*dom* x *mus*) to estimate allele-specific expression (ASE) and assign genes to eight different regulatory categories: *cis*, *trans*, *cis* X *trans*, compensatory, *cis* + *trans* opposite, *cis* + *trans* same, other, and conserved (Coolon, et al. 2014; Mack, et al. 2016).

Across all cell types and genotypes, 50-90% of genes were conserved. Comparing the two spermatogenic stages, we saw striking differences in the proportions of non-conserved genes within each regulatory category (fig. 5, supplementary table S5, Supplementary Material online). *Trans* was more common than *cis* early, whereas *trans* and *cis* made up a similar proportion of regulatory changes late (fig. 5, supplementary table S5, Supplementary Material online). Compensatory changes (compensatory and *cis*+*trans* opposite) were more common than reinforcing (*cis*+*trans* same) in both cell types, but there was a higher relative proportion of reinforcing late (fig. 5, supplementary table S5, Supplementary Material online). We focused on results for the *dom* (LEWES)♀ X *mus* (PWK)♂ cross (fig. 5) because these F1 hybrids are more fertile and therefore less likely to have misexpressed genes due to hybrid incompatibilities (Good, et al. 2010). However, the subfertile reciprocal hybrid genotype also showed similar overall proportions of genes in each regulatory category. The proportions of different regulatory mechanisms in whole testes were more similar to the late cell type (supplementary table S5, Supplementary Material online), consistent with previous studies showing high overlap in expression profiles between whole testes and spermatid stage cells (Soumillon, et al. 2013). We further verified our results using pure strain (LEWES and PWK) expression data from our phylogenetic expression dataset to determine differences in parental strain expression levels (supplementary table S5, Supplementary Material online). Finally, we evaluated the relative contributions of regulatory mechanisms contributing to expression differences between strains within each *M. musculus* subspecies using expression data from within-subspecies F1s (WSB X LEWES and CZECHII X PWK) and from the respective parental inbred strains. Consistent with the more divergent F1 hybrids, there was more *trans* than *cis* early but some variation depending on subspecies and cross-type (supplementary table S5, Supplementary Material online). In summary, early and late spermatogenesis differed in the types of regulatory mutations contributing to expression divergence, with a proportionally higher contribution of *trans*-regulatory changes early. This pattern was consistent across different degrees of evolutionary divergence and between reciprocal crosses.

**Figure 5.**
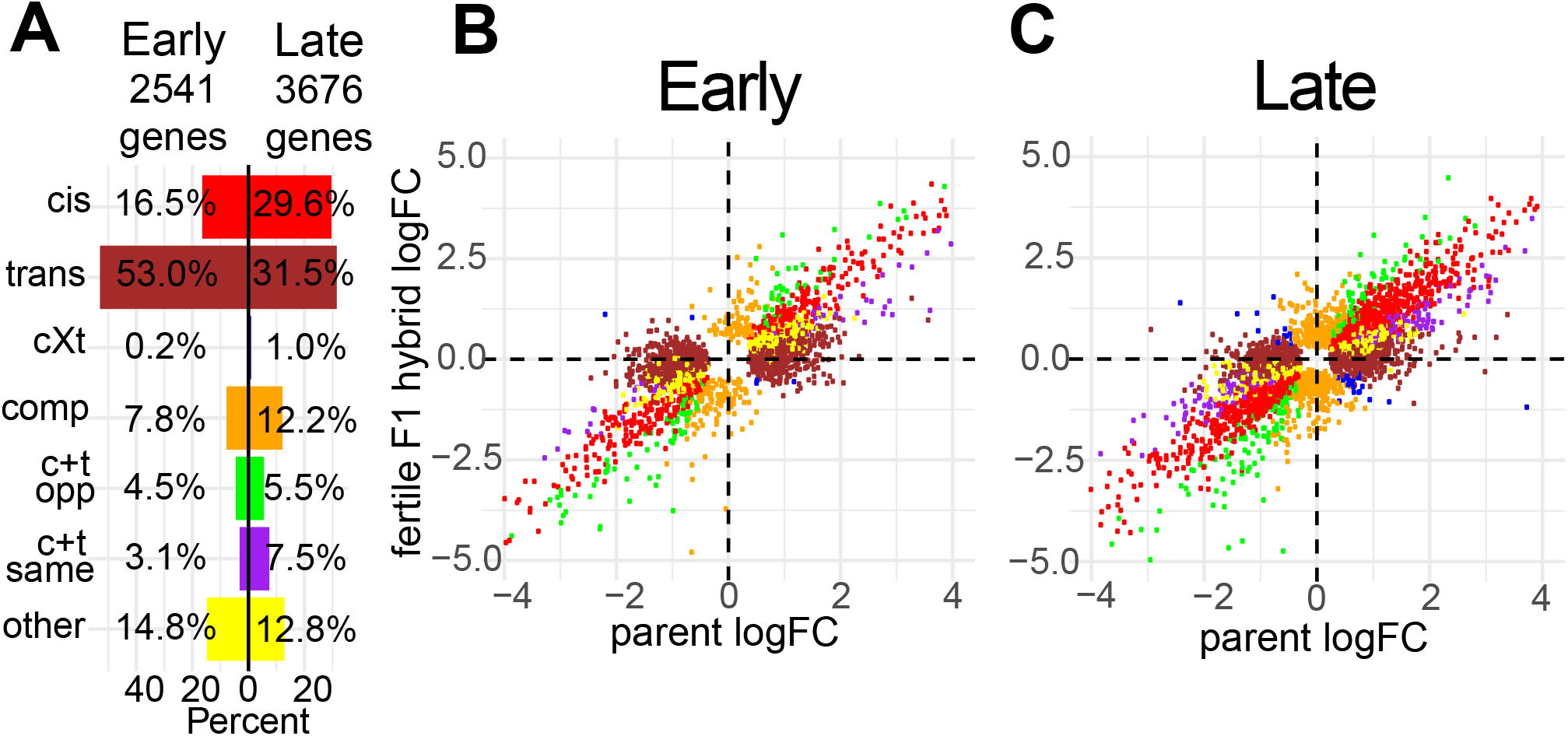
Regulatory category results for the fertile F1 hybrid (LEWES♀ X PWK♂) (*A*) Percent of non-conserved genes in each regulatory category both early and late. (*B* and *C*) Expression logFC between alleles within the fertile F1 (y-axis) plotted against the expression logFC between the parental subspecies (x-axis). Each point represents a single gene. Colors correspond to (*A*) and indicate the regulatory category to which that gene was assigned. cXt = *cis* X *trans*; comp = compensatory; c+t opp = *cis* + *trans* opposite; c+t same = *cis* + *trans* same

### *cis*-Regulatory Changes Tended to Have Larger Effects on Expression Level Divergence

Given that *trans*-regulatory changes were proportionally more common during early spermatogenesis (fig. 5), and that expression levels tended to be more conserved early (fig. 3), we hypothesized that *trans*-regulatory changes would have smaller effect sizes (Coolon, et al. 2014; Hill, et al. 2020). Consistent with this, genes with *trans* changes showed lower median divergence than those with *cis* changes (fig. 6). We saw higher divergence for reinforcing mutations based on logFC, but not EVE (fig. 6), suggesting that genes with reinforcing changes specific to the *dom* and *mus* comparison may not accumulate more divergence at deeper phylogenetic levels. For the early cell type, 26% of genes in the reinforcing category overlapped with genes that had high pairwise divergence between *dom* and *mus*, whereas only 10-16% of genes in this category overlapped with high divergence genes in other pairwise comparisons (supplementary table S6, Supplementary Material online). Similar patterns were observed for late cell type genes (supplementary table S6, Supplementary Material online). Collectively, *cis*-regulatory changes tended to have larger effects on expression divergence than *trans*-regulatory changes, and reinforcing mutations tended to have large effects on expression divergence between *mus* and *dom*, but not at deeper levels of evolutionary divergence.

**Figure 6.**
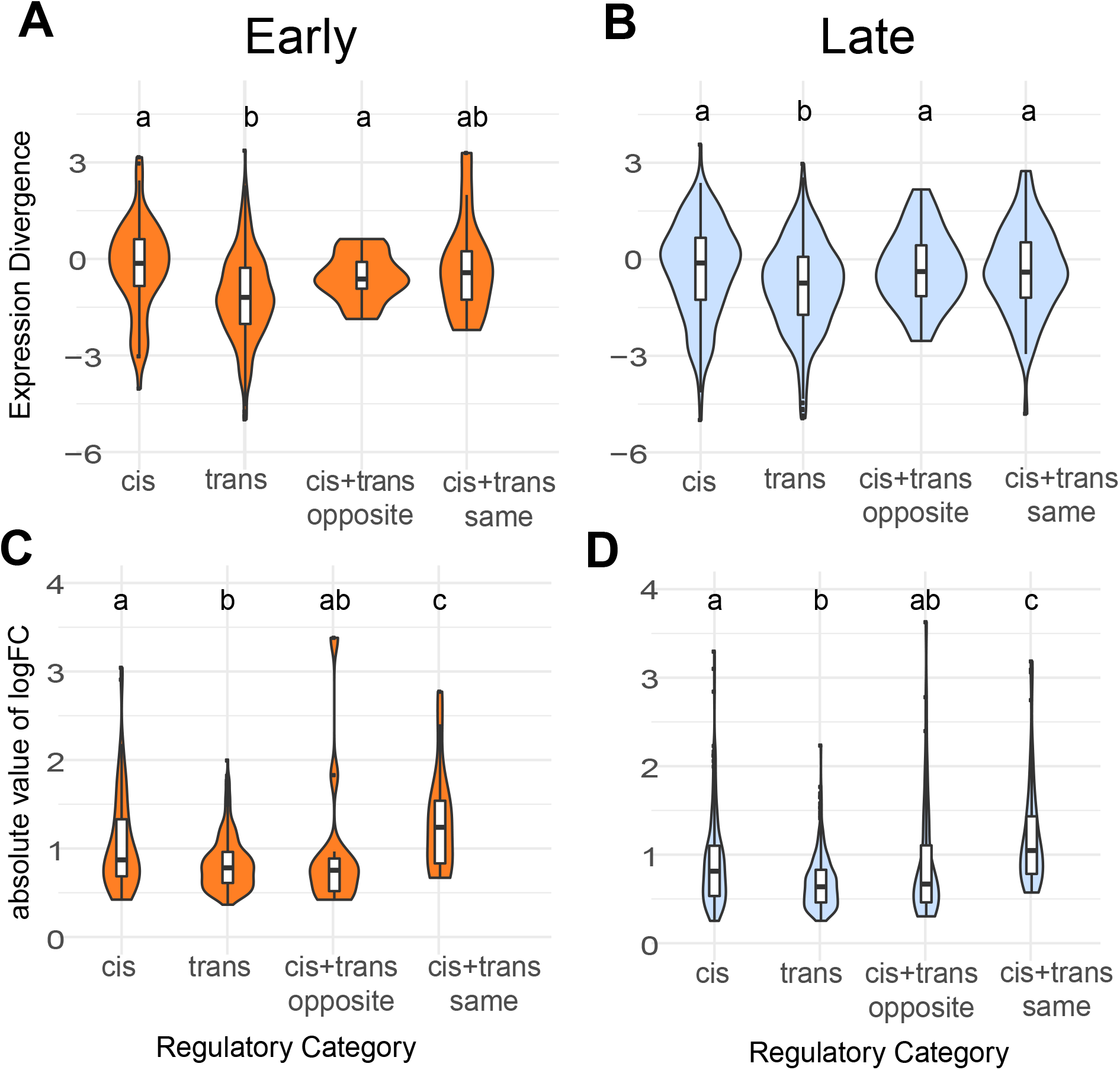
Expression divergence violin plots by regulatory category for the fertile hybrid. Expression divergence is calculated using the value from EVE (*A* and *B*) and as the absolute value of the logFC in expression between parental subspecies (*C* and *D*). Plots *A* and *C* correspond to the early cell type and plots *B* and *D* correspond to the late cell type. Letters indicate significant differences between categories based on a pairwise Wilcoxon rank sum test.

## Discussion

Developmental stage and context play an important role in shaping the molecular evolution of reproductive genes (Dean, et al. 2009; Larson, et al. 2016; Finseth and Harrison 2018; Schumacher and Herlyn 2018), with genes expressed in later developmental stages evolving more rapidly (Good and Nachman 2005; Larson, et al. 2016). However, comparing gene expression and protein divergence across developmental stages has rarely been done in a phylogenetic framework. In this study, we combined comparative genomics with cell sorting in four species to understand mouse spermatogenesis evolution across a common developmental framework. Our results give insight into how evolution proceeds at different stages of sperm development, at different molecular levels, and on different chromosome types.

### Molecular Divergence across Development

There is a long-standing prediction that early developmental stages should be more constrained, with evolutionary divergence gradually increasing across development (Abzhanov 2013), which likely contributes to more rapid molecular evolution during the later stages of sperm development. In addition, the postmeiotic stages are enriched for genes with narrower expression profiles or highly specific biological functions and are therefore expected to experience relaxed pleiotropic constraint (Eddy 2002; Good and Nachman 2005; Green, et al. 2018), also motivating our general hypothesis that the postmeiotic round spermatid stage would diverge more rapidly. Sexual selection is also likely to be a primary determinant of spermatogenic evolution, but variation in the intensity of sexual selection across spermatogenesis is not well understood (White-Cooper, et al. 2009). Sperm competition and cryptic female choice can select for changes in sperm production rate, form, or function, and many aspects of sperm morphology correlate with the intensity of post-mating sexual selection (Lüpold, et al. 2016; McLennan, et al. 2017; Pahl, et al. 2018). Rates of mitotic and initial meiotic divisions during early spermatogenesis can control the overall rate of sperm production (Ramm and Schärer 2014). Therefore, selection for increased sperm production likely acts during the development of spermatogonia, the diploid mitotic cells (White-Cooper, et al. 2009). In contrast, sexual selection shaping the form and function of mature sperm (e.g., sperm swimming speed and fertilization ability) likely acts on later developmental stages such as haploid spermatids (Alavioon, et al. 2017). However, many genes involved in mature spermatozoa functions are also highly expressed during early meiosis (da Cruz, et al. 2016), suggesting that spermatozoa may be shaped by regulatory networks operating throughout spermatogenesis.

All components of molecular evolution that we considered showed more divergence when considering genes induced in late spermatogenesis: lineage-specific expression (fig. 2), protein-coding divergence, and expression level divergence (fig. 3). On first principles, these likely result from a combination of positive selection and relaxed developmental and pleiotropic constraint (Eddy 2002; Swanson and Vacquier 2002; Winter, et al. 2004; Good and Nachman 2005; Abzhanov 2013; Green, et al. 2018). However, our study was underpowered to formally test for positive selection using likelihood ratio test approaches (Anisimova, et al. 2001; Rohlfs and Nielsen 2015). Thus, the relative contributions of positive selection and relaxed constraint to rapid spermatogenesis evolution remain unclear, especially for gene expression phenotypes.

Induced genes provided strong evidence for rapid evolution late, but results were less clear when looking at other genes. Spermatogenesis is a transcriptionally complex process, with most of the genome expressed in the testes (Soumillon, et al. 2013) and high overlap between genes expressed early and late in our dataset (table 1). For protein-coding divergence, we saw more rapid evolution late only when looking at the induced dataset, but not when looking at all expressed genes, likely because most genes in our dataset were expressed in both cell types. For expression divergence, there was more rapid evolution late even when looking at all expressed genes. This suggests that even genes with broader (i.e., non-induced) expression patterns tended to show more conserved expression early in spermatogenesis.

Testis-specific genes tended to be both induced late and rapidly evolving at the protein-coding level. Testis-specific and male-biased gene sequences often evolve rapidly, which could be the result of positive selection on genes with specific spermatogenesis functions as well as relaxed constraint because these genes tend to have highly specific functions (Meiklejohn, et al. 2003; Baines, et al. 2008; Meisel 2011; Parsch and Ellegren 2013). However, we did not see a significant faster late pattern for protein-coding or pairwise expression divergence when looking only at testis-specific genes. Although there were relatively few testis-specific genes, it appears that they tended to be rapidly evolving regardless of which spermatogenesis stage they were expressed in. If generally true, more rapid divergence late in spermatogenesis may partially reflect a higher proportion of testis-specific genes induced in the late cell type (table 1).

In addition to these broad patterns of molecular evolution, we explored the potential functional relevance of rapid divergence for specific genes (supplementary table S7, Supplementary Material online). We detected 20 genes with high (>2.5) EVE divergence in either cell type, and of these 15 were broadly expressed, but five may have specific roles in spermatogenesis (The UniProt Consortium 2020). For example, Rnf19a had an EVE value of 4.2 in the late cell type and has a known role in the formation of the sex body, which isolates the sex chromosomes in the nucleus during meiosis, a process that is required for proper spermatogenesis (Párraga and del Mazo 2000) and appears to be disrupted in sterile hybrid mice (Bhattacharyya, et al. 2013).

### Gene Expression versus Protein-Coding Divergence

Protein-coding changes alter a gene in every tissue and developmental stage in which it is expressed, whereas expression changes have the potential to be more specific (Wray, et al. 2003; Carroll 2008). Expression changes, specifically *cis*-regulatory changes, should be less constrained by pleiotropy and may underlie evolutionary changes when purifying selection acts more strongly against protein-coding divergence (Wray, et al. 2003; Carroll 2008). Under this model, we might expect to see less pronounced differences in relative expression levels when comparing early versus late stages. However, more recent work has shown that *cis*-regulatory elements such as enhancers can be highly pleiotropic, so *cis*-regulatory changes may be more constrained than once thought (Sabarís, et al. 2019; Hill, et al. 2020). If gene expression and protein-coding are subject to similar constraints, we would expect them to show similar evolutionary patterns across spermatogenesis, as we observed for autosomal genes (fig. 3).

Interestingly, despite parallel trends in relative divergence across spermatogenesis, expression level divergence and protein-coding divergence were not strongly correlated across genes, suggesting that these two types of molecular changes mostly evolve independently (Khaitovich, et al. 2005). Perhaps surprisingly, there was no overlap between genes with very rapid protein-coding divergence (dN/dS > 1.5) and high expression divergence (EVE divergence > 2.5). Likewise, only 26 genes with high pairwise expression divergence in at least one comparison (pairwise divergence metric > 1) also had high protein-coding divergence (dN/dS > 1.5; supplementary table S7, Supplementary Material online). Whether expression or protein-coding is more rapid for a particular gene may depend on factors such as expression breadth and protein function, but rarely did spermatogenic genes appear to be rapidly evolving for both gene expression and protein sequences.

We also investigated lineage-specificity. Testes and sperm tend to be enriched for lineage-specific genes (Brawand, et al. 2011) and novel genes (Schroeder, et al. 2019; Cridland, et al. 2020; Lange, et al. 2021). Lineage-specific and novel genes may be common in spermatogenesis because testes are highly transcriptionally active and have a high tissue-specific expression profile, which may allow new genes to arise without disrupting other processes (Levine, et al. 2006; Kaessmann 2010; Soumillon, et al. 2013; Zhao, et al. 2014). We found that late spermatogenesis also had proportionally more lineage-specific genes (fig. 2). Increased lineage-specificity late is consistent with and likely contributed to higher protein and expression level divergence late, as all results suggest that spermatogenesis can tolerate more genetic changes during the late stages without impacting fertility.

### X Chromosome Evolution

The X chromosome is predicted to evolve faster than the autosomes because it has a lower effective population size and is therefore more susceptible to drift, and because it is hemizygous in males so beneficial recessive mutations will fix more quickly (Charlesworth, et al. 1987; Vicoso and Charlesworth 2009). Empirical studies show evidence for a faster-X effect at the protein-coding level in many taxa, particularly for male reproductive genes (Khaitovich, et al. 2005; Baines, et al. 2008; Meisel and Connallon 2013; Parsch and Ellegren 2013; Larson, et al. 2016; Whittle, et al. 2020). Our data provide strong evidence for faster-X protein-coding evolution for both early and late spermatogenesis, demonstrating that the faster-X effect applies across genes involved in different spermatogenesis stages in mice.

Our results were more complex for expression evolution, with phylogenetic (Rohlfs and Nielsen 2015) and pairwise approaches (Meisel, et al. 2012) sometimes yielding contrasting results. In the early cell type, pairwise comparisons supported a faster-X effect, while the phylogenetic model did not (fig. 3B, supplementary table S3, Supplementary Material online). Correlations between different pairwise divergence values were relatively low on the X chromosome early, suggesting that X-linked genes with high expression level divergence in one pairwise comparison did not tend to have high divergence in other comparisons (table 2). In the late cell type, both phylogenetic and pairwise divergence metrics supported a similar rate of X-linked and autosomal expression evolution (fig. 3B, supplementary table S3, Supplementary Material online). It is well-established that lineage-specific changes can create false signatures of rapid divergence in pairwise comparisons (Felsenstein 1985), including in studies of gene expression evolution (Dunn, et al. 2018). Thus, our results highlight the importance of accounting for shared evolutionary history when inferring general evolutionary trends (Rohlfs and Nielsen 2015; Dunn, et al. 2018).

Overall, our results did not support a faster-X effect for testis gene expression evolution, in contrast to several previous studies (Khaitovich, et al. 2005; Brawand, et al. 2011; Meisel, et al. 2012). These studies were in other systems and used whole testes samples, which are made up of different cell types, so signals of expression divergence may partially reflect differences in cell type composition rather than true per cell changes in expression levels (Good, et al. 2010; Hunnicutt, et al. 2021; Yapar, et al. 2021). One previous study used cell-type specific data and found that the X chromosome showed fewer differentially expressed genes during late spermatogenesis between *mus* and *dom* (Larson, et al. 2016), and our phylogenetic sampling demonstrates that this result likely applies across mouse species.

Contrasting patterns of expression level and protein sequence evolution on the X chromosome during spermatogenesis suggest that these two levels of molecular evolution experience different evolutionary pressures (Larson, et al. 2016). Theoretical predictions for the faster-X effect on protein-coding evolution may also apply to gene expression changes, but only for *cis*-regulatory changes where the causative mutations are on the X chromosome (Meisel and Connallon 2013; Larson, et al. 2016). During spermatogenesis, the sex chromosomes undergo MSCI and PSCR, which likely imposes an overall repressive regulatory environment that constrains gene expression levels but not protein-coding changes. Disruption of MSCI and PSCR strongly impairs male fertility, so evolutionary constraints on X chromosome expression during spermatogenesis are expected to be strong (Burgoyne, et al. 2009; Good, et al. 2010; Larson, et al. 2017). Our results support the hypothesis that unique X-linked regulatory constraints reduce expression level divergence during spermatogenesis, while still allowing rapid protein-coding divergence (Larson, et al. 2016; Larson, et al. 2018a). This finding underscores how different components of molecular evolution may experience unique evolutionary pressures that result in distinct patterns of divergence (Brawand, et al. 2011; Halligan, et al. 2013; Larson, et al. 2016).

### Regulatory Mechanisms Underlying Expression Divergence

Resolving the relative contributions of *cis*-versus *trans*-acting mutations underlying expression divergence is an important step towards understanding the genetic architecture of expression phenotypes and how different evolutionary forces may act on gene expression (Benowitz, et al. 2020; Hill, et al. 2020). Although considerable progress has been made in a few key model systems on this important question (Goncalves, et al. 2012; Coolon, et al. 2014; Mack, et al. 2016; Benowitz, et al. 2020; Cridland, et al. 2020; Sánchez-Ramírez, et al. 2021), available data mostly come from whole tissues or organisms. Our results showed that the relative contribution of underlying regulatory mechanisms can differ dramatically between two cell types within a single complex tissue. While these striking differences are perhaps an expected consequence of different selective pressures acting on cellular function and developmental stage, they also underscore how difficult it is to resolve regulatory phenotypes from complex tissues.

As *cis*-regulatory changes are generally thought to be less pleiotropic (Carroll 2008; Hill, et al. 2020), we predicted that *cis*-regulatory mutations would be proportionally more common in early spermatogenesis, but we found the opposite pattern (fig. 5, supplementary table S5, Supplementary Material online). The relative contributions of *cis*- and *trans*-regulatory changes to expression divergence likely depend on other factors, including a tendency of *cis* mutations to have larger individual effect sizes (Coolon, et al. 2014; Hill, et al. 2020). We did observe proportionally more *cis*-regulatory changes of large effect during late spermatogenesis (fig. 6D) underlying higher overall expression divergence at this stage (fig. 3). Thus, differences in individual effect sizes of *cis*-versus *trans*-acting changes likely play a central role in shaping regulatory evolution across mouse spermatogenesis.

*Cis*- and *trans*-regulatory mutations can combine to affect the expression of a single gene, either in the same direction (reinforcing) or in opposite directions (compensatory; Goncalves, et al. 2012; Coolon, et al. 2014; Mack, et al. 2016). We observed a higher proportion of compensatory mutations than reinforcing mutations across both spermatogenesis cell types and in whole testes. This was expected given that gene expression tends to evolve under stabilizing selection (Rohlfs and Nielsen 2015), and it is consistent with previous studies across many tissue types in mice (Goncalves, et al. 2012; Mack, et al. 2016), flies (Coolon, et al. 2014; Benowitz, et al. 2020), and roundworms (Sánchez-Ramírez, et al. 2021). We also saw relatively more reinforcing mutations during postmeiotic spermatogenesis. Reinforcing mutations tended to have a larger effect size based on expression logFC between *mus* and *dom* (fig. 6D), thus large-effect reinforcing changes also likely contribute to higher expression level divergence in late spermatogenesis.

Given the striking differences that we saw between just two cell types, it is likely that complex tissues composed of many cell types may often give different results than isolated cell populations. Consistent with this prediction, our observed proportions of genes in each regulatory category differ from some other published results in house mouse whole tissues (i.e., liver, Goncalves, et al. 2012; whole testes, Mack, et al. 2016), primarily in that we saw a much higher proportion of genes in the *trans* category. We also found some different patterns when reanalyzing whole testes expression data from Mack, et al. (2016) that likely reflect technical differences in the analytical pipelines used between studies (supplementary table S5, see Supplementary Methods for details, Supplementary Material online). In general, our analysis used more conservative approaches to test for significant DE or ASE. Thus, only genes showing relatively pronounced differences in expression levels between genotypes or alleles were assigned to regulatory mechanisms in our study.

We also found that the relative proportion of *cis*- and *trans*-regulatory changes were similar between whole testes and the late cell type in the fertile F1 hybrid (supplementary table S5, Supplementary Material online), consistent with the observation that postmeiotic spermatids have a disproportionately large contribution to mouse whole testes expression patterns (Hunnicutt, et al. 2021). These results suggest that changes in the relative intensities of different selective pressures acting across spermatogenesis not only change the extent of expression level divergence, but also select for different mechanisms of regulatory evolution underlying these expression changes. Given this, analyzing such patterns at the level of whole organisms or tissues seems unlikely to provide a clear understanding of how mechanisms of regulatory evolution proceed in underlying cells. Indeed, even enriched cell populations as we have generated may be limited by relative purities.

By considering both expression divergence across the *Mus* phylogeny and underlying mechanisms of regulatory divergence between two lineages (*mus* and *dom*), our study also provided a novel opportunity to connect different types of regulatory changes to patterns of expression divergence at a deeper phylogenetic scale. Although *trans*-acting changes were relatively common (fig. 5), genes with *cis*-regulatory changes between *mus* and *dom* tended to have higher phylogeny-wide expression divergence than those with *trans*-regulatory changes for both cell types (fig. 6A, 6B). This suggests that genes showing *cis*-regulatory changes are also more likely to accumulate regulatory differences over time, resulting in phylogeny-wide expression divergence, whereas genes showing *trans*-regulatory changes at relatively shallow evolutionary scales tend to be relatively conserved across the *Mus* phylogeny. Genes with reinforcing changes also had relatively low phylogeny-wide expression level divergence (fig. 6A, 6B), in contrast to their high pairwise divergence between *mus* and *dom* (fig. 6C, 6D). Genes in this category likely have large-effect, lineage-specific changes in expression that may be under purifying selection over deeper phylogenetic levels. Finally, our phylogenetic contrast revealed rapid expression level divergence late in spermatogenesis. By combining these data with allele-specific expression data, we further showed that *cis*-regulatory changes are likely to underlie this rapid phylogeny-wide expression divergence in late spermatogenesis.

## Materials and Methods

### Mouse Resources

We investigated gene expression and protein-coding evolution in 12 *Mus musculus domesticus* (*dom*) individuals from four inbred strains (2 BIK/g, 3 DGA, 3 LEWES/EiJ, 4 WSB/EiJ), 8 *M. m. musculus* (*mus*) individuals from three inbred strains (2 CZII/EiJ, 3 MBS, 3 PWK/PhJ), 11 *M. spretus* (*spr*) individuals from three inbred strains (5 SEG, 2 SFM, 4 STF), and 3 *M. pahari* (*pah*) individuals from one inbred strain (3 PAHARI/EiJ; fig. 1B). By using multiple wild-derived inbred strains of *dom*, *mus*, and *spr*, we sampled natural within-species variation while also having biological replicates of genetically similar individuals. These mice were maintained in breeding colonies at the University of Montana (UM) Department of Laboratory Animal Resources (IACUC protocol 002-13). These colonies were initially established from mice purchased from The Jackson Laboratory, Bar Harbor, ME (CZECHII/EiJ, PWK/PhJ, WSB/ EiJ, LEWES/EiJ, PAHARI/EiJ) or acquired from Matthew Dean’s colonies at University of Southern California which were derived from François Bonhomme’s stocks at the University of Montpellier, Montpellier, France (MBS, BIK, DGA, STF, SFM, SEG). We weaned males at ~21 days postpartum (dpp) into same sex sibling groups and caged males individually at least 15 days prior to euthanization to avoid dominance effects on testes expression. We euthanized mice at 60-160 dpp by CO2 followed by cervical dislocation.

For expression data from reciprocal F1 males, we used FACS enriched expression data from (Larson, et al. 2017). These data include males from reciprocal F1 crosses between different inbred strains within each *M. musculus* subspecies (*mus*: CZECHII females X PWK males, *dom*: WSB females X LEWES males), as well as reciprocal *mus* and *dom* F1 hybrids (LEWES females X PWK males and PWK females X LEWES males), allowing us to compare results at two different levels of divergence (i.e., within and between lineages). We also analyzed whole testes expression data from Mack, et al. (2016) to compare FACS enriched cell-types to whole testes, including crosses between different strains within each *M. musculus* subspecies (LEWES females X WSB males and PWK females X CZII males) and the same reciprocal F1 hybrid crosses to those in (Larson, et al. 2017).

### Testis Cell Sorting and RNAseq

We collected testes from mice immediately following euthanization and isolated cells at different stages of spermatogenesis using Fluorescence Activated Cell Sorting (FACS; Getun, et al. 2011). The full FACS protocol is available on GitHub (https://github.com/goodest-goodlab/good-protocols/tree/main/protocols/FACS). Briefly, we decapsulated testes and washed them twice with 1mg/mL collagenase (Worthington Biochemical), 0.004mg/ mL DNase I (Qiagen), and GBSS (Sigma), followed by disassociation with 1mg/mL trypsin (Worthington Biochemical) and 0.004mg/mL DNase I. We then inactivated trypsin with 0.16mg/mL fetal calf serum (Sigma).

For each wash and disassociation step, we incubated and agitated samples at 33°C for 15 minutes on a VWR minishaker at 120 rpm. We stained cells with 0.36mg/mL Hoechst 33324 (Invitrogen) and 0.002mg/mL propidium iodide, filtered with a 40μm cell filter, and sorted using a FACSAria IIu cell sorter (BD Biosciences) at the UM Center for Environmental Health Sciences Fluorescence Cytometry Core. We periodically added 0.004mg/mL DNase I as needed during sorting to prevent DNA clumps from clogging the sorter. We sorted cells into 15μL beta-mercaptoethanol (Sigma) per 1mL of RLT lysis buffer (Qiagen) and kept samples on ice whenever they were not in the incubator or the cell sorter. For this study, we focused on two cell populations: early meiotic spermatocytes (leptotene/ zygotene) and postmeiotic round spermatids. We extracted RNA using the Qiagen RNeasy Blood and Tissue Kit and checked RNA integrity with a Bioanalyzer 2000 (Agilent) or TapeStation 2200 (Agilent). All samples except one had RIN ≥ 7 (supplementary table S8, Supplementary Material online). We prepared RNAseq libraries using the Agilent SureSelect protocol and sequenced samples at the Hudson Alpha Institute for Biotechnology using Illumina NextSeq (75bp single end).

### *Mus* strain phylogeny

We generated the phylogeny in fig. 1B using available exome (Chang, et al. 2017; Sarver, et al. 2017) and whole genome (Keane, et al. 2011; Thybert, et al. 2018) sequence data (PRJNA326865, PRJNA323493, PRJEB2003, PRJEB14896). Genotypes were based on iterative mapping assemblies relative to the house mouse reference genome (mm10) conducted using pseudo-it v3.0 (Sarver, et al. 2017; https://github.com/goodest-goodlab/pseudo-it/tree/kopania-etal-2021) that restricts genotyping to targeted exons. We ran pseudo-it with one iteration to generate consensus fasta files for each sample. We then extracted exons, aligned these regions using MAFFT v7.271 (Katoh and Standley 2013), converted to PHYLIP format using AMAS (Borowiec 2016), and inferred a maximum likelihood concatenated tree using IQ-TREE v2.1.4-beta (Nguyen, et al. 2014).

### Processing and Quantification of Gene Expression Data

We used R version 3.6.3 and Bioconductor version 3.10 for all analyses. We trimmed raw reads for adaptors and low-quality bases using expHTS (Streett, et al. 2015) and mapped trimmed reads with TopHat version 2.1.0 (Kim, et al. 2013). Genome assemblies were previously published for all four lineages (Keane, et al. 2011; Thybert, et al. 2018), allowing us to map reads to the correct assembly and reduce reference bias (Sarver, et al. 2017). Mapping rates were consistent across lineages (supplementary table S8, Supplementary Material online). To select orthologous genes among the four lineages, we used BiomaRt (Durinck, et al. 2005; Durinck, et al. 2009) to identify one-to-one Ensembl orthologs and retained only those that were present in all genome assemblies and the mouse reference build GRCm38.

We used edgeR 3.28.1 (Robinson, et al. 2010) to normalize expression data, calculate fragments per kilobase per million reads (FPKM), and perform differential expression (DE) analyses. A gene was defined as “expressed” in our dataset if it had an FPKM > 1 in at least eight samples. A gene was expressed in a particular lineage and cell type if it had an FPKM > 1 in all samples of that lineage and cell type. A gene was considered induced in a particular cell type if its median FPKM in that cell type across all lineages was greater than two times its median FPKM in the other cell type across all lineages. Testis-specific genes were those only expressed in testis based on the mouse tissue expression data from (Chalmel, et al. 2007).

We defined lineage-specific genes in two ways. First, we used a log fold change (logFC) method in which a gene was considered lineage-specific if its median expression level in a lineage was greater than two times its median expression level in any of the other three lineages. Second, we used a Bayesian approach to determine if a gene is active or inactive in an expression dataset based on transcript levels as implemented with the program Zigzag (Thompson, et al. 2020). Genes identified as being active (posterior P > 0.5) in one lineage and inactive (posterior P < 0.5) in the other lineages were considered lineage-specific. We ran Zigzag twice and only included genes with consistent active or inactive assignments between the two runs. Both the logFC and Zigzag analyses were performed for each cell type, so a gene could be lineage-specific in one cell type but not the other. For each lineage, we determined the proportion of expressed (logFC) or active (Zigzag) genes that were lineage-specific and used a Pearson’s χ2 test to determine if one cell type had greater lineage-specificity than the other.

### Protein-Coding Divergence

We used the “iqtree-omp” command in IQTree version 1.5.5 (Nguyen, et al. 2014) to infer a mouse species tree based on gene trees estimated from the reference sequences for all four mouse lineages (Keane, et al. 2011; Thybert, et al. 2018). We took the longest transcript for all one-to-one orthologs and aligned these using MAFFT v7.271 (Katoh and Standley 2013) and converted to PHYLIP format using AMAS (Borowiec 2016). We used a custom script to exclude genes that did not begin with a start codon, had early stop codons, or had sequence lengths that were not multiples of three. We then used the Codeml program in the PAML package to calculate protein-coding divergence and test for positive selection on protein-coding genes (Yang 2007). We used the M0 model to calculate phylogeny-wide dN/dS for each gene, which we report as the overall protein-coding divergence values. We also performed a likelihood ratio test between the M8 and M8a site-based models to test for positive directional selection on each gene (Swanson, et al. 2003).

### Differential Expression

We performed all analyses of expression level divergence for three different gene sets: expressed genes, induced genes, and testis-specific genes. To calculate expression divergence in a phylogenetic framework, we used the EVE model (Rohlfs and Nielsen 2015), which performs a phylogenetic ANOVA using an Ornstein-Uhlenbeck model to evaluate divergence while controlling for evolutionary relatedness. We report expression divergence from EVE as −log(*beta*), where *beta* is a metric from EVE that represents the ratio of within-lineage variance to between-lineage evolutionary divergence. By taking the negative log, higher positive numbers correspond to greater evolutionary divergence. We excluded genes with extremely low divergence values [−log(*beta_i_*)<−5] because this subset did not show a linear relationship between evolutionary divergence and population variance and therefore violated underlying assumptions of the EVE model (supplementary fig. S6, Supplementary Material online).

We also calculated expression divergence in a pairwise framework (Meisel, et al. 2012). This method takes the difference in expression level between two lineages and normalizes based on the average expression of the gene in both lineages:

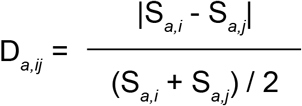

Da,ij is the divergence of gene a between lineages i and j. Sa,i is the median FPKM of gene a in lineage i, and Sa,j is the median FPKM of gene a in lineage j. We also calculated the logFC in expression between every pairwise comparison of lineages as an additional pairwise divergence metric (Robinson, et al. 2010). For the EVE, pairwise divergence, and logFC methods, we compared relative expression divergence between cell types and between the X chromosome and autosomes using a Wilcoxon rank-sum test. We tested if certain cell types or chromosome types showed greater correlation among pairwise divergence values using Spearman’s rank correlation.

We also compared relative expression divergence on the X chromosome versus the autosomes using the proportion of DE genes on each chromosome (Good, et al. 2010; Larson, et al. 2016). First, we calculated the proportion of expressed genes that are DE across all autosomes. We then multiplied this proportion by the number of genes expressed on each chromosome to calculate the expected number of DE genes for each chromosome. We plotted the observed number of DE genes against the expected number and used a hypergeometric test to evaluate if each chromosome is over- or under-enriched for DE genes.

### Allele-Specific Expression and Regulatory Divergence

We used the modtools and lapels-suspenders pipelines (Huang, et al. 2014) to reduce mapping bias and to assign the parental origin of reads in F1 individuals (See Supplementary Methods for details, Supplementary Material online). This approach requires mapping to pseudogenomes generated using modtools to resolve differences in genome coordinates between different references. We used published pseudogenomes for WSB and PWK, which incorporate single nucleotide variants (SNVs) and indels from these strains into the GRCm38 mouse reference build (Huang, et al. 2014). For LEWES and CZECHII, we generated our own pseudogenomes with modtools version 1.0.2 using published VCF files (Morgan, et al. 2016; Larson, et al. 2018b). We developed a custom pipeline (See Supplementary Methods for details, Supplementary Material online) to assign genes to regulatory categories following previous recommendations (Coolon, et al. 2014; Mack, et al. 2016; Combs and Fraser 2018; Benowitz, et al. 2020). To determine significant differences between cell types, we performed a Pearson’s χ2 test followed by false discovery rate correction for multiple tests.

## Data Availability

RNAseq data generated for this project are available through the National Center for Biotechnology Information under accession PRJNA735780. Individual sample accessions are in supplementary table S8. A table of genes in our analyses and whether they were considered expressed, induced, or active in each cell type is available in supplementary table S9.

## Author contributions

J.M.G conceived and funded the project. E.E.K.K., E.L.L., and J.M.G. designed the experiments. E.L.L. and S.K. performed the mouse husbandry and breeding experiments. C.C., S.K., and E.L.L. performed mouse dissections and cell sorts. C.C. and E.L.L. prepared the sequencing libraries. E.E.K.K. analyzed the data. E.E.K.K., E.L.L., and J.M.G. wrote the manuscript with input from all authors.

## Acknowledgements

We would like to thank Pamela K. Shaw and the UM Fluorescence Cytometry Core supported by an Institutional Development Award from the NIGMS (P30GM103338), the UM Genomics Core supported by the M.J. Murdock Charitable Trust, the UM Lab Animal Resources staff, Gregg Thomas for assistance generating the phylogeny in fig. 1B, Nathanael Herrera for mouse photos, and the Good Lab for helpful advice. This work was supported by grants from the Eunice Kennedy Shriver National Institute of Child Health and Human Development of the National Institutes of Health (R01-HD073439, R01-HD094787 to JMG). EEKK was supported by the National Science Foundation Graduate Research Fellowship Program (DGE-1313190). ELL was supported by the National Science Foundation (DEB-2012041). Any opinions, findings, and conclusions or recommendations expressed in this material are those of the author(s) and do not necessarily reflect the views of the National Science Foundation or the National Institutes of Health.

