## Supplementary_Material for "Molecular Evolution across Mouse Spermatogenesis"

### Supplementary Methods

**Regulatory categories:** Because the publicly available PWK pseudogenome is based on a VCF that is much older than the CZECHII VCF, the median quality scores are much higher in the CZECHII VCF which causes mapping bias when mapping CZECHII X PWK individuals. To address this, we called variants from publicly available PWK sequence reads using the Genome Analysis Tool Kit (GATK) version 4.1.7.0 program HaplotypeCaller. We also downsampled from the CZECHII sequence reads to match the read counts available for PWK and generated a new CZECHII VCF using HaplotypeCaller. We used modtools to generate new pseudogenomes for PWK and CZECHII based on these modified VCF files and only used these pseudogenomes for mapping CZECHII X PWK individuals.

We trimmed raw reads using trimmomatic version 0.35 (Bolger, et al. 2014) and mapped reads using TopHat v2.1.1 (Kim, et al. 2013), consistent with (Mack, et al. 2016). After mapping reads to both parent pseudogenomes, we converted coordinates to match the mouse reference build GRCm38 using lapels (pylapels version 0.2.0 in Lapels 1.1.1). We then used suspenders (pysuspenders version 0.2.5 in Suspenders 0.2.5), which merges lapels outputs from mappings to the two different parents and assigns reads to either parent based on sequence variants and mapping quality scores. The *mus* and *dom* subspecies are closely related and have a $D_{xy}$ of about 0.5% (Geraldes, et al. 2008), which means most reads will map equally well to both parents. For F1 hybrids, we excluded reads that mapped equally well to both parents. For parents, we mapped reads to both pseudogenomes and only kept reads that were assigned to the correct parent. For example, we would map a PWK sample to both

PWK and LEWES, run it through the full lapels-suspenders pipeline, and only keep reads that were assigned to PWK. This ensured that both F1 and parent data were treated the same and removed genes that were DE between the parents but could not be evaluated for ASE due to a lack of variants. Because these data included a combination of paired-end and single-end data, we removed the second read from all pairs in which both reads mapped, and then converted all SAM flags to single end flags. We then downsampled reads from these bam files such that both parents had a similar number of reads to those assigned to each F1 allele (Coolon, et al. 2014; Mack, et al. 2016). This gave us similar power to detect DE and ASE. We counted the number of reads mapping to each gene with HTSeq-count (Anders, et al. 2014). (supplementary fig. S7)

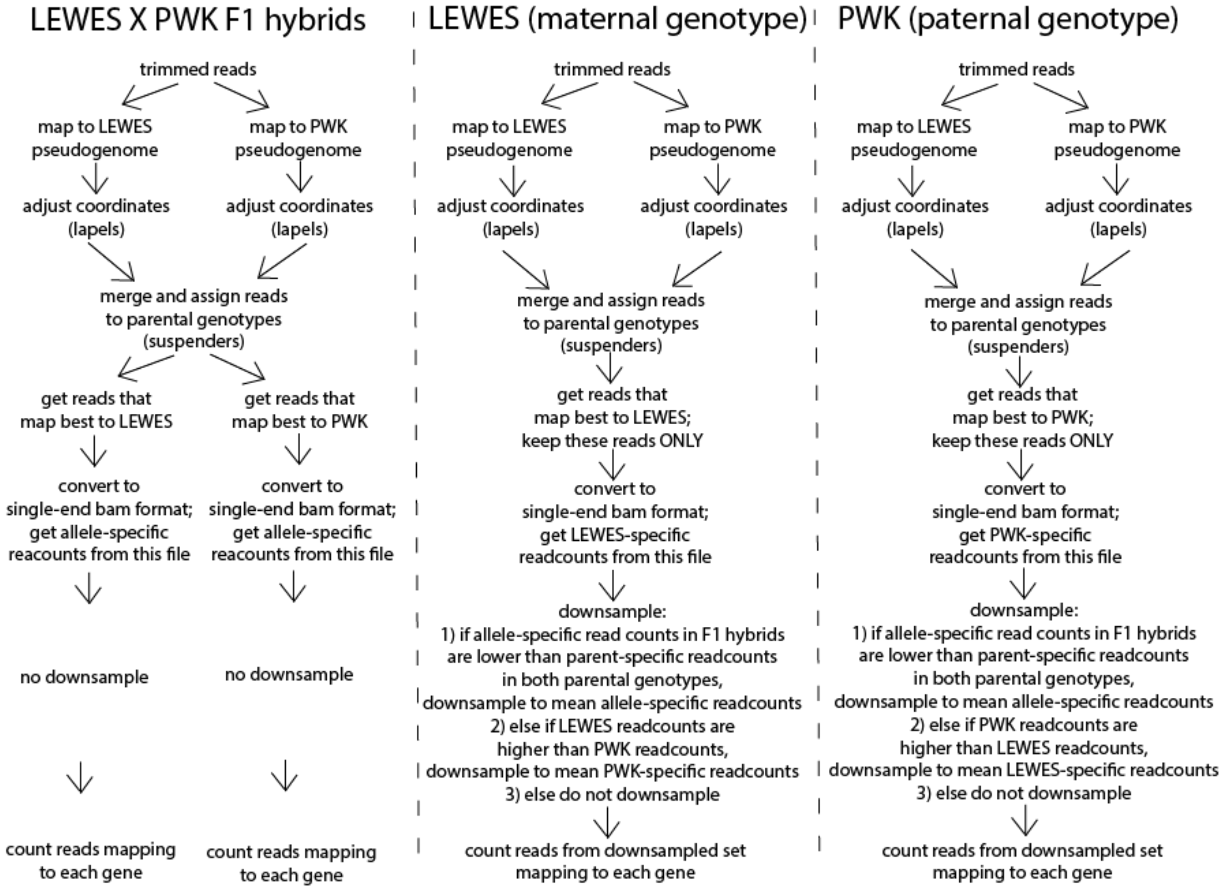


**Fig. S7.** An example of the read mapping, lapels-suspenders, and downsampling pipeline used to assess regulatory categories.

Our custom pipeline to assign genes to general regulatory categories as defined in previous studies (Coolon, et al. 2014; Mack, et al. 2016; Combs and Fraser 2018; Benowitz, et al. 2020) was as follows. First, we used a negative binomial test implemented in edgeR to determine if a gene is DE between the parental genotypes. We then also used the edgeR negative binomial model to test if the gene showed ASE within the F1 hybrids, similar to previous studies (Combs and Fraser 2018; Benowitz, et al. 2020). We used a Fisher's Exact Test (FET) to determine if the expression difference between the parents was significantly different from the allelic expression difference within the F1 hybrids (Coolon, et al. 2014; Mack, et al. 2016; Benowitz, et al. 2020). If a gene was not DE between the parental genotypes and showed no evidence for ASE in the F1 hybrids, it was treated as conserved. If a gene was not DE, but had ASE with a significant FET, it was assigned to the compensatory category. Genes that were DE but had no ASE with a significant FET were assigned to the *trans* category. A gene with both DE and ASE and a non-significant FET was considered regulated in *cis*. Genes with both DE and ASE plus a significant FET presumably had some combination of *cis* and *trans* mutations acting on gene expression. We further broke this category down in the following ways. Genes were assigned to the *cis* X *trans* category if DE and ASE were in opposite directions (e.g., the gene was more highly expressed in the maternal parent, but the paternal allele was more highly expressed in the F1 hybrid). Genes were assigned to the *cis* + *trans* same category if DE and ASE were in the same direction and the expression difference was greater between the parental genotypes. This category represents reinforcing mutations, or *cis* and *trans* acting in the same direction. Genes were assigned to the *cis* + *trans* opposite category if DE and ASE were in the same direction and the expression difference was greater in the F1 hybrid. This category represents weak compensatory mutations, where *cis* and *trans* are acting in opposite directions but have not fully compensated each other because the gene is still DE between the parent lineages. (supplementary fig. S8)

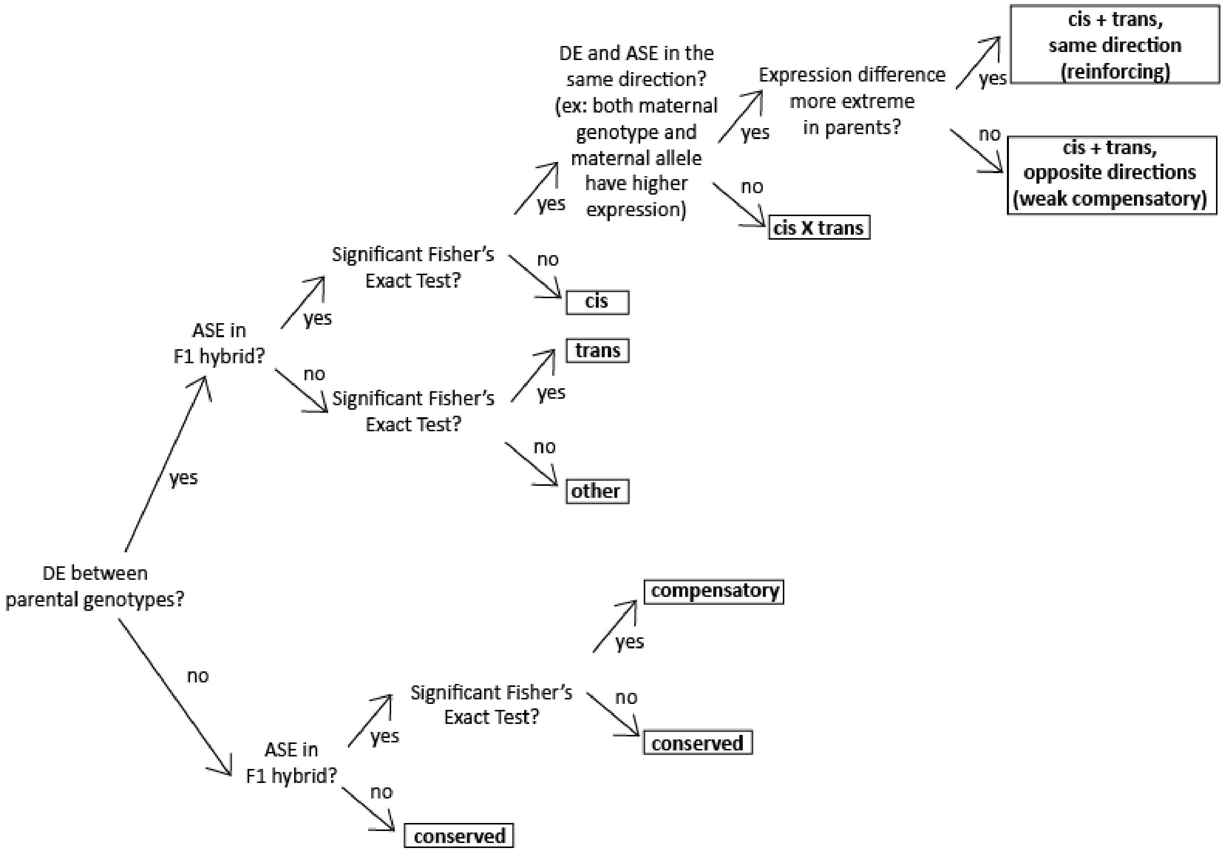


**Fig. S8.** Flow chart showing the pipeline used to assign genes to regulatory categories.

For each F1 cross, we used expression data from the parental inbred strains as the "parent" in our regulatory category analyses. For the hybrid F1s, we repeated the analyses using data from within-subspecies F1s as the "parent" to compare the effects of having a homozygous parent vs a heterozygous parent. For example, we performed the analyses with LEWES♀ X PWK♂ F1 hybrids using LEWES and PWK as the parents, and then repeated the analysis using data from the same F1 hybrids and using data from WSB♀ X LEWES♂ and CZECHII♀ X PWK♂ F1 mice as the parents. We report results based on within-subspecies F1 parents in the main text, and report all results in (supplementary table S5).

**Reanalysis of whole-testis data from Mack, et al. (2016):** Our pipeline assigned many more genes to the *trans* category and many fewer genes to the *cis* X *trans* category than reported previously (Mack, et al. 2016; supplementary table S5). To investigate the reason for this potential inconsistency, we also reanalyzed these whole testes data using a binomial test following Mack, et al. (2016), which should be more sensitive (i.e., less conservative) than a negative binomial approach for detecting expression differences. The binomial test gave results more similar to those reported by Mack, et al. (2016), with proportionally fewer genes assigned to the *trans* category and a higher proportion of genes in the *cis* X *trans* category.

We also explored gene-level differences in category assignment between the two approaches and found large groups of genes that were assigned to one category using the binomial approach that were then consistently assigned to a different category using the negative binomial approach (supplementary table S10). For example, many genes assigned to *cis+trans* (same direction) using the binomial test were categorized as *trans* using the negative binomial test. These relatively subtle differences make sense because genes with significant expression divergence between parents and less extreme expression divergence between alleles in the F1s will be assigned to one of these two categories. The key distinction is that genes in the *trans* category do not show a significant difference in ASE, and therefore, our more conservative method for considering alleles to be differentially expressed in F1s will likely assign more genes to the *trans* category (supplementary fig. S8). Other common changes in category assignment between the binomial and negative binomial approaches are consistent with a more conservative method for calling genes DE or ASE (supplementary table S10, supplementary fig. S8).

However, we note that our more conservative analytical method likely explains most, but not all, of the quantitative differences between our results and those from (Mack, et al. 2016). Other inconsistencies likely result from differences in how F1 sequencing reads were bioinformatically assigned to parents. Although the general conceptual frameworks were similar, Mack, et al. (2016) used a custom script to assign reads, while we used the lapels and suspenders pipeline from modtools (Holt, et al. 2013; Huang, et al. 2014). The number of reads assigned to each parent were similar across both studies for most samples, but there were some notable differences and individual sample outliers that may have contributed to differences in regulatory category assignment for some genes (supplementary table S11).

**Supplementary Figures**

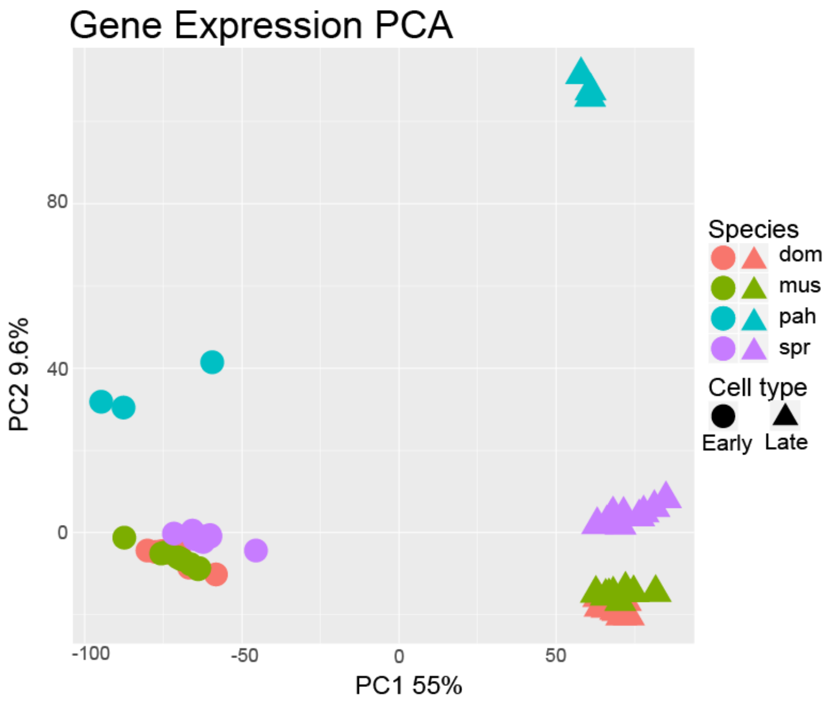


**Fig. S1.** Principal component analysis based on expression levels of genes expressed in either cell type. Circles represent the early cell type and triangles represent the late cell type. Colors correspond to different lineages. Cell type explains most of the variance (PC1, 55%) followed by lineage (PC2, 9.6%).

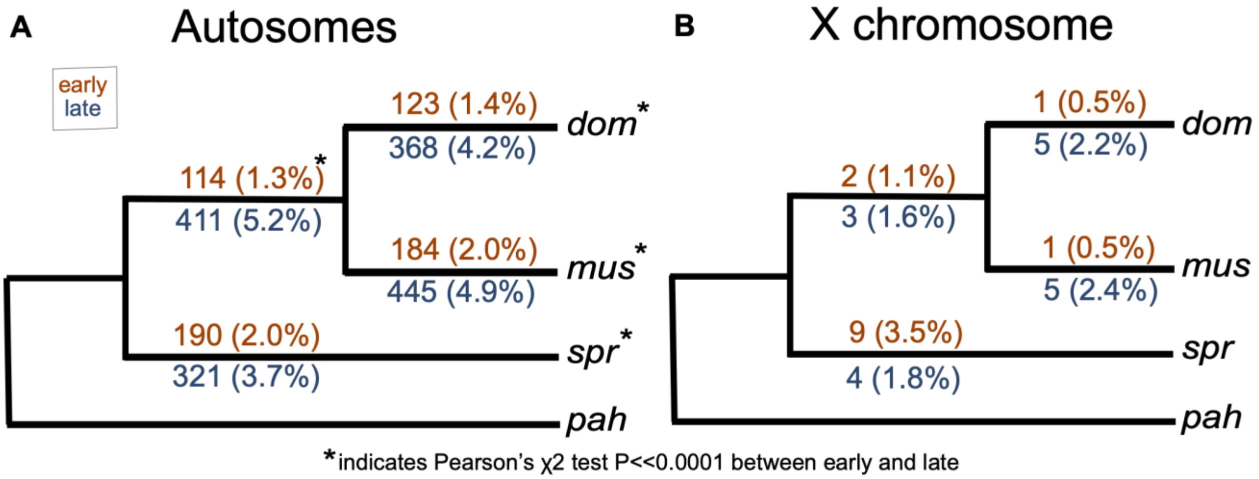


**Fig. S2.** Number of genes that are lineage-specific on each internal branch of the mouse phylogeny used in this study based on a logFC approach. Numbers in parentheses are the percent of active genes that are lineage-specific. Results are presented separately for the autosomes (*A*) and X chromosome (*B*). Orange values above each branch represent the early cell type and blue values below represent the late cell type. Asterisks indicate a significant difference between early and late on that branch based on a Pearson's $\chi^2$ test.

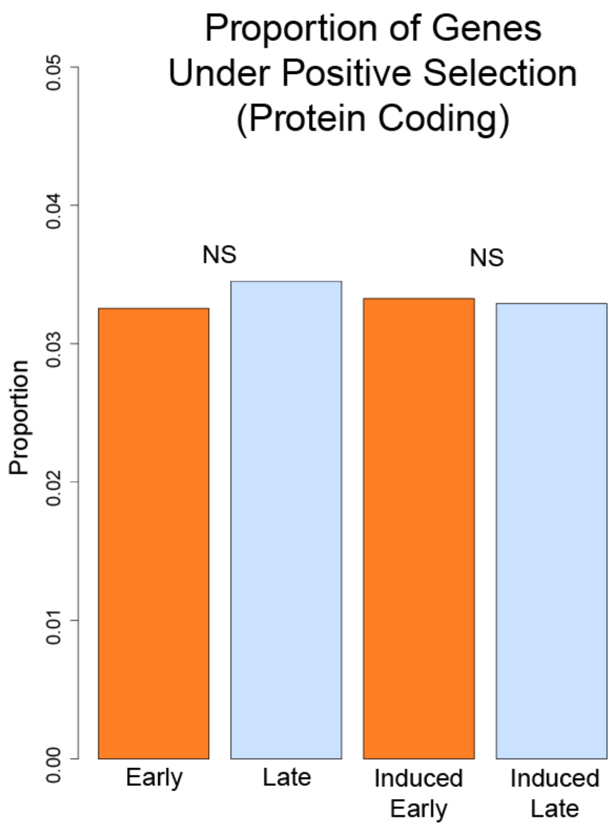


**Fig. S3.** Proportion of genes expressed or induced in each cell type under positive selection at the protein-coding level. NS = not significantly different (Pearson's $\chi^2$ P > 0.05 after FDR correction)

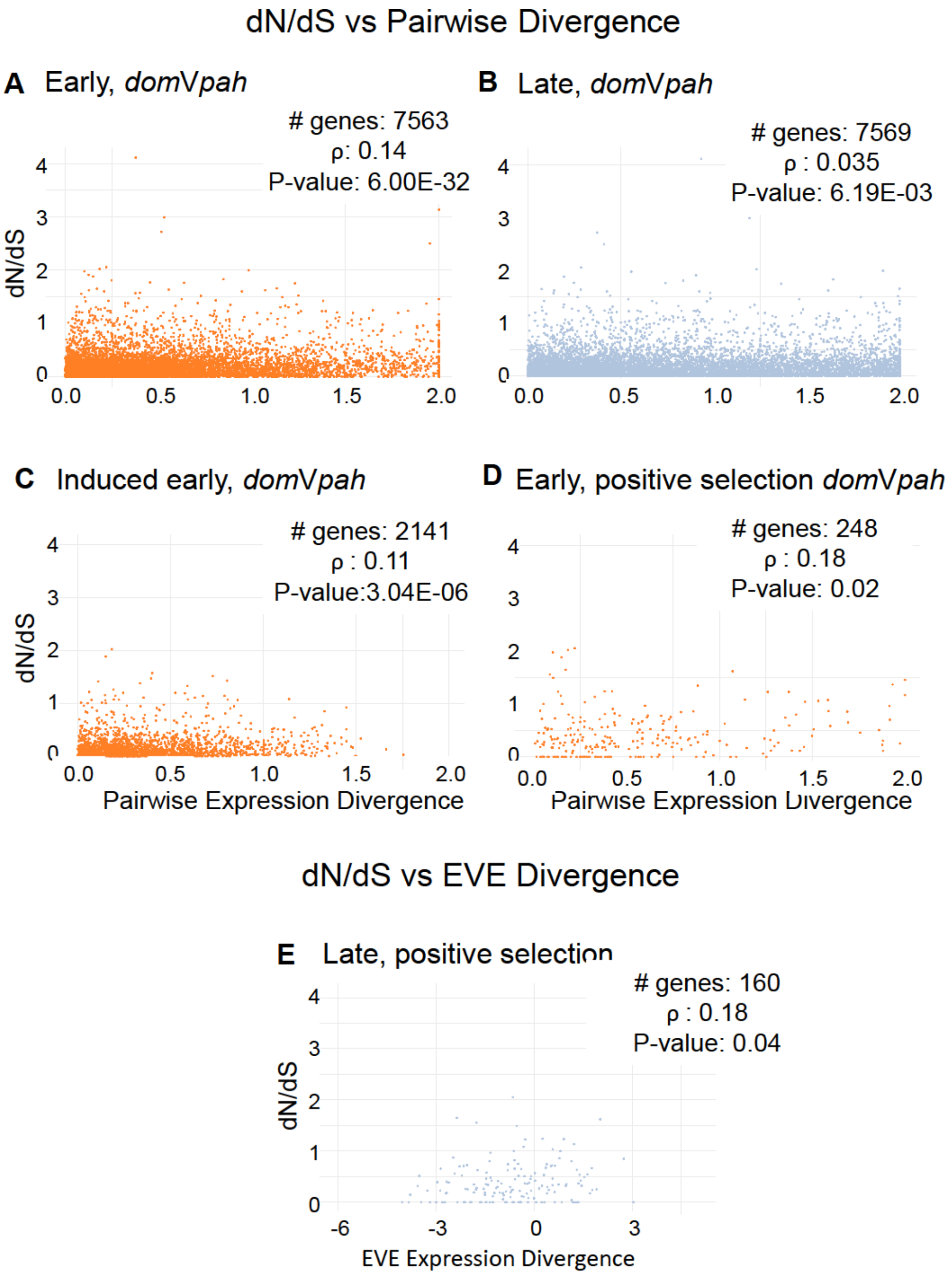


**Fig. S4.** Some examples showing weak positive correlations between protein-coding and expression level divergence. We chose to show the *dom* vs *pah* comparisons for the pairwise divergence plots, but other pairwise comparisons show similar patterns. See supplementary table S5 for Spearman's ρ and p-values for all comparisons. (*A*-*D*) dN/dS vs pairwise expression divergence for: (*A*) all genes expressed early, (*B*) all genes expressed late, (*C*) genes induced early, (*D*) genes expressed early and under positive selection for protein-coding; (*E*) dN/dS vs EVE expression divergence for all genes expressed late that are under positive selection for protein-coding.

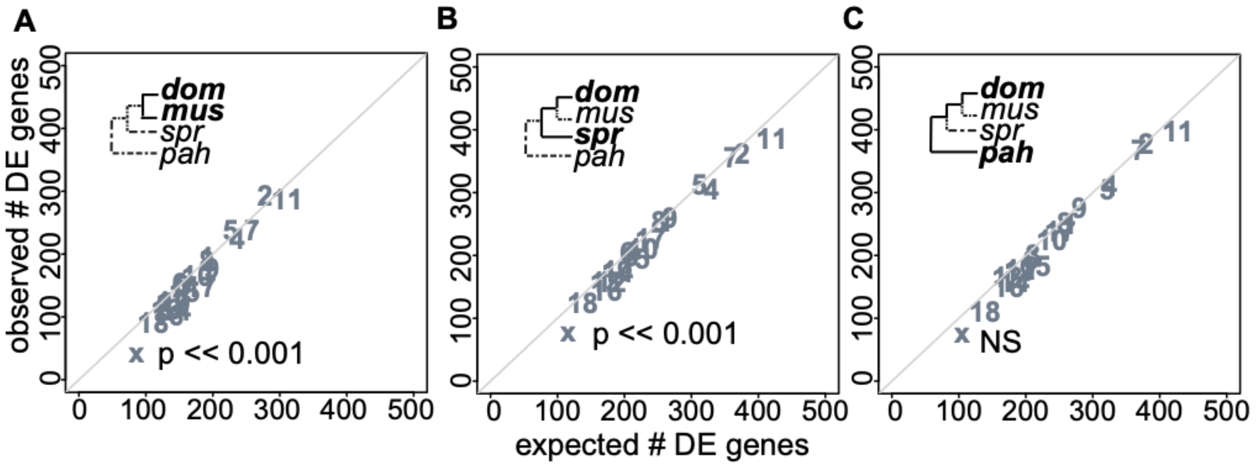


**Fig. S5.** Observed versus expected number of genes differentially expressed (DE) in late spermatogenesis for three pairwise comparisons at different levels of evolutionary divergence: (*A*) *dom* versus *mus*, (*B*) *spr* versus *dom,* and (*C*) *pah* versus *dom*. Each point represents a different chromosome. The diagonal line is the one-to-one line at which the observed number of DE genes equals the expected number. P-values are shown for the X chromosome only. They are based on a hypergeometric test for enrichment and corrected for multiple tests using a false discovery rate correction. A significant p-value indicates that the observed number of DE genes is different from the expected number.

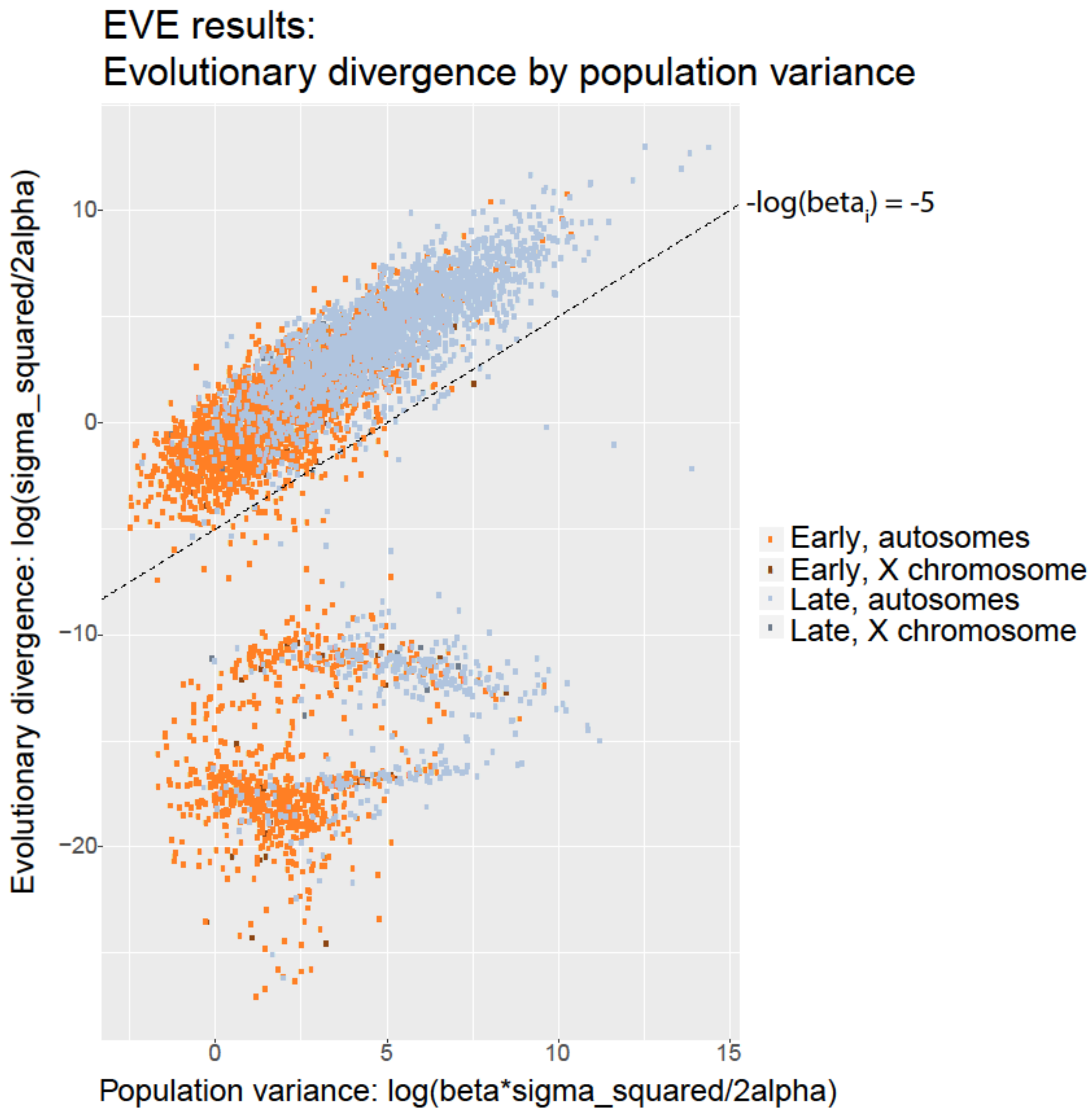


**Fig. S6.** EVE model evolutionary (between lineage) variance plotted against population (within lineage) variance in expression level. The EVE model assumes a linear relationship between these two values. Because genes with a divergence value [-log($beta_i$)] below $-5$ violated this assumption, we excluded them from our analyses of expression divergence (below dashed line). This figure is based on fig. 1 from Rohlfs and Nielsen (2015). Each point represents a single gene, and points are colored by cell type and chromosome type (lighter for autosome or darker for X chromosome).

### Supplementary Tables

**Table S1.** Number of genes and median dN/dS values for genes expressed, induced, testis-specific, or testis-specific and induced at different spermatogenesis stages and different chromosomes.

| | Autosomes early | | X early | | Autosomes late | | X late | |
|---|---|---|---|---|---|---|---|---|
| | n | dN/dS | n | dN/dS | n | dN/dS | n | dN/dS |
| **Expressed** | 5729 | 0.131 | 167 | 0.182 | 5462 | 0.136 | 124 | 0.375 |
| **Induced** | 2046 | 0.105 | 54 | 0.251 | 1711 | 0.201 | 61 | 0.411 |
| **Testis-specific (TS)** | 350 | 0.282 | 16 | 0.587 | 424 | 0.297 | 24 | 0.579 |
| **TS and induced** | 32 | 0.259 | 6 | 0.745 | 329 | 0.306 | 19 | 0.543 |

**Table S2.** P-values for Wilcoxon Rank Sum tests comparing median dN/dS values between different groups of genes. Dark shaded boxes provide evidence for greater divergence in late spermatogenesis, and light shaded boxes provide evidence for faster X evolution.

| | Early vs Late, Autosomes | Early vs Late, X Chromosome | X vs Autosomes, Early | X vs Autosomes, Late |
|---|---|---|---|---|
| **Expressed** | 0.16399 | 0.00034 | 0.0061 | 7.10E-14 |
| **Induced** | <2.00E-16 | 0.0488 | 0.00015 | 1.40E-07 |
| **Testis-specific (TS)** | 1 | 1 | 0.0015 | 5.10E-06 |
| **TS and Induced** | 0.2103 | 0.1009 | 0.0013 | 0.0013 |

**Table S3.** Summary of gene expression divergence results using different methods to quantify expression divergence. Each row is a different method or set of genes, and each column is a different comparison between either cell types or chromosome types. "None" means no significant difference. For pairwise comparisons, comparisons that had a result different from the general trend are in parentheses.

| | **Early vs Late, Autosomes** | **Early vs Late, X Chromosome** | **X vs Autosomes, Early** | **X vs Autosomes, Late** |
|---|---|---|---|---|
| **EVE, all genes** | faster late | faster late | slower-X | none |
| **EVE, induced genes** | faster late | faster late | slower-X | none |
| **EVE, testis-specific genes** | faster late | none | none | none |
| **pairwise divergence, all genes** | faster late | faster late (slower late domVspr) | faster-X (none domVpah and slower-X musVspr) | none (faster-X domVmus and musVpah, slower-X domVspr) |
| **pairwise divergence, induced genes** | faster late | none (slower late domVspr) | faster-X (none musVspr) | none |
| **pairwise divergence, testis-specific genes** | none | none | none | none (faster-X domVpah, musVpah, sprVpah) |
| **logFC, all genes** | faster late (none musVdom) | faster late (slower late domVspr) | faster-X (none musVspr) | faster-X (none domVspr and slower-X musVspr) |
| **logFC, induced genes** | faster late | faster late (slower late domVmus, domVspr, and sprVpah) | faster-X (none musVspr) | faster-X (none domVmus and musVspr) |
| **logFC, testis-specific genes** | slower late (faster late domVmus, none domVpah) | none (slower late domVmus) | none (faster-X domVmus) | none |
| **proportion DE genes, all genes** | NA | NA | none (slower-X musVspr and domVpah) | slower-X (none sprVpah) |

| | | | | |
|---|---|---|---|---|
| **proportion DE genes, induced genes** | NA | NA | none (slower-X musVspr) | none (slower-X domVmus) |
| **proportion DE genes, testis specific** | NA | NA | none | none |

**Table S4.** Relationship between protein-coding and expression level divergence. Rows in bold are significant based on Spearman's rank correlation. P-values are adjusted using an FDR correction for multiple tests. PW = pairwise expression divergence; ED = EVE phylogeny-wide expression divergence

| Divergence comparison | Species comparison | Cell type | Induced? | Positive selection (PAML)? | No. of genes | ρ | P-value |
|---|---|---|---|---|---|---|---|
| **dN/dS vs PW** | **domVmus** | **early** | **no** | **no** | **7562** | **0.13** | **5.19E-27** |
| **dN/dS vs PW** | **domVspr** | **early** | **no** | **no** | **7569** | **0.16** | **3.67E-44** |
| **dN/dS vs PW** | **musVspr** | **early** | **no** | **no** | **7569** | **0.17** | **6.23E-50** |
| **dN/dS vs PW** | **domVpah** | **early** | **no** | **no** | **7563** | **0.14** | **6.00E-32** |
| **dN/dS vs PW** | **musVpah** | **early** | **no** | **no** | **7561** | **0.13** | **1.97E-29** |
| **dN/dS vs PW** | **sprVpah** | **early** | **no** | **no** | **7567** | **0.13** | **7.96E-28** |
| **dN/dS vs PW** | **domVmus** | **late** | **no** | **no** | **7570** | **0.03** | **2.17E-02** |
| **dN/dS vs PW** | **domVspr** | **late** | **no** | **no** | **7571** | **0.04** | **6.56E-04** |
| **dN/dS vs PW** | **musVspr** | **late** | **no** | **no** | **7569** | **0.05** | **2.27E-04** |
| **dN/dS vs PW** | **domVpah** | **late** | **no** | **no** | **7569** | **0.04** | **6.19E-03** |
| **dN/dS vs PW** | **musVpah** | **late** | **no** | **no** | **7562** | **0.03** | **7.23E-03** |
| **dN/dS vs PW** | **sprVpah** | **late** | **no** | **no** | **7562** | **0.05** | **2.95E-04** |
| **dN/dS vs PW** | **domVmus** | **early** | **yes** | **no** | **2141** | **0.07** | **6.19E-03** |
| **dN/dS vs PW** | **domVspr** | **early** | **yes** | **no** | **2141** | **0.09** | **2.49E-04** |
| **dN/dS vs PW** | **musVspr** | **early** | **yes** | **no** | **2141** | **0.09** | **2.49E-04** |
| **dN/dS vs PW** | **domVpah** | **early** | **yes** | **no** | **2141** | **0.11** | **3.04E-06** |
| **dN/dS vs PW** | **musVpah** | **early** | **yes** | **no** | **2141** | **0.08** | **6.97E-04** |
| **dN/dS vs PW** | **sprVpah** | **early** | **yes** | **no** | **2141** | **0.09** | **7.75E-05** |
| dN/dS vs PW | domVmus | late | yes | no | 1760 | 0.02 | 0.57 |
| dN/dS vs PW | domVspr | late | yes | no | 1760 | 0.01 | 0.70 |
| dN/dS vs PW | musVspr | late | yes | no | 1760 | 0.01 | 0.87 |
| dN/dS vs PW | domVpah | late | yes | no | 1760 | 0.00 | 0.92 |
| dN/dS vs PW | musVpah | late | yes | no | 1760 | -0.01 | 0.70 |
| dN/dS vs PW | sprVpah | late | yes | no | 1760 | -0.04 | 0.23 |
| dN/dS vs PW | domVmus | early | no | yes | 248 | 0.09 | 0.31 |
| dN/dS vs PW | domVspr | early | no | yes | 250 | 0.12 | 0.10 |

| | | | | | | | |
|---|---|---|---|---|---|---|---|
| **dN/dS vs PW** | **musVspr** | **early** | **no** | **yes** | **249** | **0.20** | **0.00** |
| **dN/dS vs PW** | **domVpah** | **early** | **no** | **yes** | **248** | **0.18** | **0.02** |
| **dN/dS vs PW** | **musVpah** | **early** | **no** | **yes** | **247** | **0.19** | **0.00** |
| **dN/dS vs PW** | **sprVpah** | **early** | **no** | **yes** | **248** | **0.21** | **0.00** |
| dN/dS vs PW | domVmus | late | no | yes | 250 | 0.08 | 0.32 |
| **dN/dS vs PW** | **domVspr** | **late** | **no** | **yes** | **250** | **0.17** | **0.02** |
| **dN/dS vs PW** | **musVspr** | **late** | **no** | **yes** | **249** | **0.18** | **0.00** |
| dN/dS vs PW | domVpah | late | no | yes | 250 | 0.06 | 0.45 |
| dN/dS vs PW | musVpah | late | no | yes | 249 | 0.06 | 0.48 |
| dN/dS vs PW | sprVpah | late | no | yes | 250 | 0.09 | 0.25 |
| dN/dS vs PW | domVmus | early | yes | yes | 67 | 0.19 | 0.24 |
| dN/dS vs PW | domVspr | early | yes | yes | 67 | 0.00 | 1.00 |
| dN/dS vs PW | musVspr | early | yes | yes | 67 | 0.16 | 0.31 |
| dN/dS vs PW | domVpah | early | yes | yes | 67 | 0.08 | 0.60 |
| dN/dS vs PW | musVpah | early | yes | yes | 67 | -0.02 | 0.90 |
| dN/dS vs PW | sprVpah | early | yes | yes | 67 | 0.09 | 0.60 |
| dN/dS vs PW | domVmus | late | yes | yes | 57 | -0.15 | 0.39 |
| dN/dS vs PW | domVspr | late | yes | yes | 57 | -0.21 | 0.23 |
| dN/dS vs PW | musVspr | late | yes | yes | 57 | -0.04 | 0.84 |
| dN/dS vs PW | domVpah | late | yes | yes | 57 | -0.10 | 0.58 |
| dN/dS vs PW | musVpah | late | yes | yes | 57 | -0.10 | 0.57 |
| dN/dS vs PW | sprVpah | late | yes | yes | 57 | -0.16 | 0.36 |
| dN/dS vs ED | NA | early | no | no | 4473 | 0.03 | 0.10 |
| dN/dS vs ED | NA | late | no | no | 4755 | 0.02 | 0.43 |
| dN/dS vs ED | NA | early | yes | no | 1544 | 0.02 | 0.56 |
| dN/dS vs ED | NA | late | yes | no | 1490 | -0.01 | 0.89 |
| dN/dS vs ED | NA | early | no | yes | 144 | -0.11 | 0.31 |
| **dN/dS vs ED** | **NA** | **late** | **no** | **yes** | **160** | **0.18** | **0.04** |
| dN/dS vs ED | NA | early | yes | yes | 55 | -0.12 | 0.51 |
| dN/dS vs ED | NA | late | yes | yes | 48 | 0 | 1.00 |

**Table S5.** Proportion of genes in each regulatory category. P-values are based on a Pearson's chi-squared test for differences between the early and late cell types after FDR correction for multiple tests. The first column shows the results presented in Mack, et al. (2016). For the first two columns, genes in the "conserved" category are grouped into the "other" category for direct comparison with results from Mack, et al. (2016). *cXt =cisXtrans;* comp = compensatory; c+*t,* opp = *cis + trans,* opposite; *c+t,* same = *cis + trans,* same

| | **fertile F1 hybrid, intra-subspecific F1 parents (binomial test)*** | | **fertile F1 hybrid, intra-subspecific F1 parents** | | | | **fertile F1 hybrid, pure strain parents** | | | **sterile F1 hybrid, intra-subspecific F1 parents** | | | | **sterile F1 hybrid, pure strain parents** | | | ***dom* only** | | | ***mus* only** | | |
|---|---|---|---|---|---|---|---|---|---|---|---|---|---|---|---|---|---|---|---|---|---|---|
| | **Mack et al. 2016** | **re-analysis (Modtools)** | **whole testes** | **early** | **late** | **p-val** | **early** | **late** | **p-val** | **whole testes** | **early** | **late** | **p-val** | **early** | **late** | **p-val** | **early** | **late** | **p-val** | **early** | **late** | **p-val** |
| ***cis*** | 24% | 21% | 22% | 17% | 30% | 9.57E-47 | 11% | 33% | 4.86E-157 | 14% | 18% | 30% | 6.59E-34 | 12% | 31% | 3.17E-145 | 14% | 22% | 5.42E-07 | 8% | 12% | 1.33E-04 |
| ***trans*** | 9% | 17% | 36% | 53% | 32% | 2.46E-119 | 59% | 28% | 0.00E+00 | 34% | 51% | 33% | 1.39E-84 | 57% | 29% | 3.31E-315 | 46% | 29% | 1.01E-27 | 59% | 28% | 1.14E-130 |
| ***cXt*** | 7% | 10% | 0% | 0% | 1% | 1.08E-04 | 1% | 2% | 4.34E-04 | 0% | 0% | 1% | 6.60E-03 | 1% | 2% | 1.33E-02 | 0% | 3% | 5.14E-05 | 3% | 4% | 7.63E-01 |
| **comp** | 13% | 8% | 16% | 8% | 12% | 2.65E-01 | 7% | 10% | 5.62E-05 | 14% | 10% | 12% | 1.94E-03 | 9% | 10% | 8.45E-04 | 12% | 9% | 9.94E-03 | 12% | 18% | 3.97E-07 |
| ***c+t*, opp** | 16% | 9% | 12% | 5% | 6% | 3.02E-02 | 2% | 3% | 2.88E-05 | 15% | 4% | 5% | 3.17E-02 | 2% | 4% | 1.13E-09 | 3% | 6% | 5.55E-04 | 3% | 9% | 3.54E-12 |
| ***c+t*, same** | 8% | 17% | 11% | 3% | 8% | 1.20E-16 | 5% | 9% | 2.50E-14 | 13% | 4% | 7% | 1.26E-09 | 7% | 10% | 4.72E-10 | 10% | 11% | 8.86E-01 | 7% | 16% | 2.23E-19 |
| **other** | 23% | 18% | 4% | 15% | 13% | 3.28E-03 | 14% | 15% | 3.88E-01 | 11% | 13% | 13% | 5.78E-01 | 13% | 15% | 2.12E-03 | 14% | 21% | 8.07E-07 | 8% | 15% | 3.58E-10 |
| **Total # genes** | 9851 | 9478 | 1430 | 2541 | 3676 | NA | 3291 | 4067 | NA | 1129 | 2416 | 3258 | NA | 3820 | 3859 | NA | 910 | 1768 | NA | 1215 | 1643 | NA |

*includes conserved genes in "other" category

**Table S6.** Proportion of genes in each regulatory category that showed high pairwise expression divergence. High pairwise divergence is defined as genes in the top 25% of divergence values for a given pairwise comparison. Each row represents a different pairwise comparison and cell type. Highlighted boxes represent the proportion of genes in the *cis* + *trans* same (reinforcing) category that also were highly divergent between *dom* and *mus*. Of all genes in the reinforcing category, a higher proportion overlap with *dom* vs *mus* highly divergent genes than with genes highly divergent in other pairwise comparisons.

| **Early** | ***cis*** | ***trans*** | ***cis* X *trans*** | **compensatory** | ***cis + trans* opposite** | ***cis + trans* same** | **other** |
|---|---|---|---|---|---|---|---|
| *dom*V*mus* | 0.21 | 0.137 | 0 | 0.151 | 0.261 | 0.263 | 0.183 |
| *dom*V*spr* | 0.126 | 0.092 | 0 | 0.131 | 0.113 | 0.125 | 0.151 |
| *mus*V*spr* | 0.103 | 0.107 | 0 | 0.146 | 0.122 | 0.15 | 0.146 |
| *dom*V*pah* | 0.093 | 0.108 | 0.2 | 0.126 | 0.087 | 0.113 | 0.143 |
| *mus*V*pah* | 0.105 | 0.106 | 0.2 | 0.151 | 0.13 | 0.163 | 0.149 |
| *spr*V*pah* | 0.081 | 0.103 | 0.2 | 0.131 | 0.113 | 0.1 | 0.141 |
| **Late** | ***cis*** | ***trans*** | ***cis* X *trans*** | **compensatory** | ***cis + trans* opposite** | ***cis + trans* same** | **other** |
| *dom*V*mus* | 0.198 | 0.055 | 0.029 | 0.031 | 0.129 | 0.215 | 0.07 |
| *dom*V*spr* | 0.106 | 0.08 | 0.143 | 0.092 | 0.163 | 0.142 | 0.11 |
| *mus*V*spr* | 0.122 | 0.073 | 0.171 | 0.067 | 0.158 | 0.102 | 0.085 |
| *dom*V*pah* | 0.128 | 0.078 | 0.229 | 0.098 | 0.144 | 0.109 | 0.104 |
| *mus*V*pah* | 0.108 | 0.083 | 0.229 | 0.101 | 0.124 | 0.117 | 0.089 |
| *spr*V*pah* | 0.107 | 0.091 | 0.114 | 0.116 | 0.099 | 0.095 | 0.1 |

**Table S7.** Genes with evidence for rapid evolution during spermatogenesis (protein-coding, phylogeny-wide expression, or pairwise expression) that may also have testis-biased expression. Comparisons with high pairwise divergence have a divergence value > 1. Chr = chromosome; LZ = leptotene/zygotene spermatocytes ("early"); RS = round spermatids ("late"); EVE = -log(*beta*$_i$) value from the EVE model

| Gene ID | Gene Name | Chr | dN/dS | Positive Selection (PAML)? | Induced ? | High Pairwise Expression Divergence (LZ) | High Pairwise Expression Divergence (RS) | EVE (LZ) | EVE (RS) | Expression Pattern |
|---|---|---|---|---|---|---|---|---|---|---|
| ENSMUSG00000022280 | Rnf19a | 15 | 0.14 | no | yes, in RS | none | none | -0.266 | 4.159 | primarily testis |
| ENSMUSG00000022602 | Arc | 15 | 0 | no | no | none | none | 0.023 | 4.251 | primarily testis and brain |
| ENSMUSG00000056209 | Npm3 | 19 | 0.12 | no | no | none | none | 3.388 | 2.687 | primarily testis |
| ENSMUSG00000037101 | Ttc29 | 8 | 2.05 | yes | yes, in RS | none | none | -1.134 | -0.643 | primarily testis |
| ENSMUSG00000049761 | Pmis2 | 7 | 1.55 | yes | yes, in RS | none | none | -1.773 | -1.76 | testis-specific |
| ENSMUSG00000027317 | Ppp1r14d | 2 | 2.71 | no | NA | domVmus, musVspr | domVmus, domVspr, musVspr, musVpah, sprVpah | NA | NA | has a testis-specific isoform |

**Table S8.** RNAseq metadata for each sample.

| Lineage | Strain | Sample Name | Cell Type | Cell Sort Date | Mouse Age at Sort Date | RNA concentration after cell sort RNA extraction (ng/µL) | RIN | # Raw Reads | # Mapped Reads | SRA Accession |
|---|---|---|---|---|---|---|---|---|---|---|
| *dom* | BIK/g | BIK_4665.1M_LZ | LZ | 10/17/2013 | 95 | 8.5 | 9.6 | 28574278 | 15792758 | SAMN19597717 |
| *dom* | BIK/g | BIK_4665.1M_RS | RS | 10/17/2013 | 95 | 4.2 | 8.5 | 38477703 | 29776396 | SAMN19597718 |
| *dom* | BIK/g | BIK_4665.2M_LZ | LZ | 10/18/2013 | 96 | 12.4 | 9.7 | 11156170 | 9679873 | SAMN19597719 |
| *dom* | BIK/g | BIK_4665.2M_RS | RS | 10/18/2013 | 96 | 3.4 | 8.1 | 38039743 | 30916495 | SAMN19597720 |
| *dom* | DGA | DGA_5406.1M_LZ | LZ | 10/3/2013 | 94 | 8.2 | 9.8 | 24007340 | 14415796 | SAMN19597721 |
| *dom* | DGA | DGA_5406.1M_RS | RS | 10/3/2013 | 94 | 4.7 | 8.7 | 10757881 | 9926211 | SAMN19597722 |
| *dom* | DGA | DGA_5406.2M_LZ | LZ | 10/10/2013 | 101 | 7.4 | 9.7 | 16740461 | 14956442 | SAMN19597723 |
| *dom* | DGA | DGA_5406.2M_RS | RS | 10/10/2013 | 101 | 7.7 | 8.6 | 10080556 | 9278016 | SAMN19597724 |
| *dom* | DGA | DGA_5406.3M_LZ | LZ | 10/11/2013 | 102 | 10.6 | 9.6 | 26873371 | 22540332 | SAMN19597725 |
| *dom* | DGA | DGA_5406.3M_RS | RS | 10/11/2013 | 102 | 9.7 | 8.4 | 50306765 | 45439841 | SAMN19597726 |
| *dom* | LEWES/EiJ | LL.LL125.1M.LZ | LZ | 6/4/2015 | 80 | 14.2 | 9.7 | 41523441 | 32816986 | SAMN19597727 |
| *dom* | LEWES/EiJ | LL.LL125.1M.RS | RS | 6/4/2015 | 80 | 17.6 | 7.9 | 32174124 | 28997946 | SAMN19597728 |
| *dom* | LEWES/EiJ | LL.LL125.2M.LZ | LZ | 6/2/2015 | 78 | 6.1 | 8.8 | 38838198 | 34572862 | SAMN19597729 |
| *dom* | LEWES/EiJ | LL.LL125.2M.RS | RS | 6/2/2015 | 78 | 5.6 | 8.5 | 16433187 | 15281121 | SAMN19597730 |
| *dom* | LEWES/EiJ | LL.LL125.3M.LZ | LZ | 6/1/2015 | 77 | 7.4 | 9.5 | 25464043 | 23141321 | SAMN19597731 |
| *dom* | LEWES/EiJ | LL.LL125.3M.RS | RS | 6/1/2015 | 77 | 5.6 | 8.6 | 32759463 | 29987663 | SAMN19597732 |
| *dom* | WSB/EiJ | WW.WW87.2M_LZ | LZ | 5/13/2014 | 66 | 12.2 | 9.6 | 1783976 | 1570753 | SAMN19597733 |
| *dom* | WSB/EiJ | WW.WW87.2M_RS | RS | 5/13/2014 | 66 | 4 | 8.6 | 15238105 | 14266130 | SAMN19597734 |
| *dom* | WSB/EiJ | WW.WW87.3M_LZ | LZ | 5/14/2014 | 67 | 10.6 | 9.6 | 18188548 | 16621233 | SAMN19597735 |

| | | | | | | | | | | |
|---|---|---|---|---|---|---|---|---|---|---|
| *dom* | WSB/EiJ | WW.WW87.3M_RS | RS | 5/14/2014 | 67 | 7.7 | 8.6 | 11897490 | 11086144 | SAMN19597736 |
| *dom* | WSB/EiJ | WW.WW87.4M_LZ | LZ | 5/30/2014 | 83 | 9.8 | 9.7 | 9943638 | 9062201 | SAMN19597737 |
| *dom* | WSB/EiJ | WW.WW87.4M_RS | RS | 5/30/2014 | 83 | 3.1 | 8.7 | 13971859 | 12895786 | SAMN19597738 |
| *dom* | WSB/EiJ | WW.WW89.8M_LZ | LZ | 4/29/2014 | 61 | 6.4 | 9.4 | 11641744 | 10377128 | SAMN19597739 |
| *dom* | WSB/EiJ | WW.WW89.8M_RS | RS | 4/29/2014 | 61 | 4.3 | 8.3 | 18384101 | 16980022 | SAMN19597740 |
| *mus* | CZECHII/EiJ | CC.CC153.4M.LZ | LZ | 5/20/2015 | 87 | 6.5 | 9.2 | 36237077 | 24720082 | SAMN19597741 |
| *mus* | CZECHII/EiJ | CC.CC153.4M.RS | RS | 5/20/2015 | 87 | 3.3 | 8.2 | 8995430 | 8273438 | SAMN19597742 |
| *mus* | CZECHII/EiJ | CC.CC153.5M.LZ | LZ | 5/22/2015 | 89 | 8 | 8.8 | 24808708 | 18264983 | SAMN19597743 |
| *mus* | CZECHII/EiJ | CC.CC153.5M.RS | RS | 5/22/2015 | 89 | 7.5 | 7.3 | 39256981 | 32049212 | SAMN19597744 |
| *mus* | MBS | MBS_4527.1M_LZ | LZ | 9/20/2013 | 63 | 6.8 | 9.5 | 22446803 | 20502488 | SAMN19597745 |
| *mus* | MBS | MBS_4527.1M_RS | RS | 9/20/2013 | 63 | 10.9 | 8.2 | 12665610 | 11785311 | SAMN19597746 |
| *mus* | MBS | MBS_4527.2M_LZ | LZ | 9/26/2013 | 69 | 25.9 | 8.8 | 18148591 | 16500321 | SAMN19597747 |
| *mus* | MBS | MBS_4527.2M_RS | RS | 9/26/2013 | 69 | 10.5 | 8.2 | 24450942 | 22406598 | SAMN19597748 |
| *mus* | MBS | MBS_4527.3M_LZ | LZ | 10/2/2013 | 75 | 18.3 | 9.5 | 26762674 | 24061812 | SAMN19597749 |
| *mus* | MBS | MBS_4527.3M_RS | RS | 10/2/2013 | 75 | 6.9 | 8.3 | 15396056 | 14315196 | SAMN19597750 |
| *mus* | PWK/PhJ | PP.PP.98.4M.LZ | LZ | 6/18/2015 | 73 | 35 | 9 | 11808242 | 10876393 | SAMN19597751 |
| *mus* | PWK/PhJ | PP.PP.98.4M.RS | RS | 6/18/2015 | 73 | 13.8 | 8.6 | 72582309 | 67215275 | SAMN19597752 |
| *mus* | PWK/PhJ | PP.PP98.3M.LZ | LZ | 6/17/2015 | 72 | 35.3 | 8.5 | 7372674 | 6519250 | SAMN19597753 |
| *mus* | PWK/PhJ | PP.PP98.3M.RS | RS | 6/17/2015 | 72 | 13.8 | 8.1 | 19931551 | 18654440 | SAMN19597754 |
| *mus* | PWK/PhJ | PP.PP98.5M.LZ | LZ | 6/24/2015 | 79 | 15.5 | 9.5 | 11848083 | 10572199 | SAMN19597755 |
| *mus* | PWK/PhJ | PP.PP98.5M.RS | RS | 6/24/2015 | 79 | 6.8 | 8.7 | 16044407 | 15062711 | SAMN19597756 |
| *spr* | SEG | SEG_4130_LZ | LZ | 3/5/2013 | 66 | 6 | 9.4 | 25968746 | 23559901 | SAMN19597757 |
| *spr* | SEG | SEG_4130_RS | RS | 3/5/2013 | 66 | 5.2 | 8.3 | 25844180 | 23258335 | SAMN19597758 |

| | | | | | | | | | | |
|---|---|---|---|---|---|---|---|---|---|---|
| *spr* | SEG | SEG_4156_LZ | LZ | 9/19/2013 | 130 | 12.4 | 8.6 | 20360141 | 17686631 | SAMN19597759 |
| *spr* | SEG | SEG_4156_RS | RS | 9/19/2013 | 130 | 4 | 7.6 | 22932882 | 20960724 | SAMN19597760 |
| *spr* | SEG | SEG_4176_LZ | LZ | 2/26/2013 | 100 | 4.2 | 8.9 | 22777686 | 20079778 | SAMN19597761 |
| *spr* | SEG | SEG_4176_RS | RS | 2/26/2013 | 100 | 6.8 | 7.2 | 14405903 | 13311983 | SAMN19597762 |
| *spr* | SEG | SEG_4197_LZ | LZ | 3/11/2013 | 72 | 14 | 9 | 23944237 | 20931833 | SAMN19597763 |
| *spr* | SEG | SEG_4197_RS | RS | 3/11/2013 | 72 | 5.7 | 8.4 | 10443918 | 9663280 | SAMN19597764 |
| *spr* | SEG | SEG_4700.1M_LZ | LZ | 9/9/2013 | 148 | 16.7 | 8.8 | 13571853 | 12253820 | SAMN19597765 |
| *spr* | SEG | SEG_4700.1M_RS | RS | 9/9/2013 | 148 | 6.1 | 7.8 | 24534179 | 22456069 | SAMN19597766 |
| *spr* | SFM | SFM_4513_LZ | LZ | 3/7/2013 | 74 | 17.3 | 8.6 | 13616782 | 12154832 | SAMN19597767 |
| *spr* | SFM | SFM_4513_RS | RS | 3/7/2013 | 74 | 7.4 | 8.4 | 6887627 | 6355085 | SAMN19597768 |
| *spr* | SFM | SFM_4514_LZ | LZ | 3/12/2013 | 79 | 7.6 | 9 | 16693417 | 14970665 | SAMN19597769 |
| *spr* | SFM | SFM_4514_RS | RS | 3/12/2013 | 79 | 11.3 | 7.8 | 13874143 | 12800911 | SAMN19597770 |
| *spr* | STF | STF_4495.1M_LZ | LZ | 9/11/2013 | 150 | 13.7 | 9 | 7705953 | 7049222 | SAMN19597771 |
| *spr* | STF | STF_4495.1M_RS | RS | 9/11/2013 | 150 | 4.3 | 7.9 | 22281038 | 20273895 | SAMN19597772 |
| *spr* | STF | STF_4515_LZ | LZ | 3/20/2013 | 84 | 4.2 | 9.1 | 14293633 | 13197893 | SAMN19597773 |
| *spr* | STF | STF_4515_RS | RS | 3/20/2013 | 84 | 1.8 | 9.3 | 8019452 | 7356732 | SAMN19597774 |
| *spr* | STF | STF_4516_LZ | LZ | 3/4/2013 | 68 | 4.6 | 9.8 | 17387559 | 15715819 | SAMN19597775 |
| *spr* | STF | STF_4516_RS | RS | 3/4/2013 | 68 | 6 | 8.8 | 15006751 | 13932584 | SAMN19597776 |
| *spr* | STF | STF_4517_LZ | LZ | 3/8/2013 | 72 | 20.2 | 9.2 | 32586439 | 29128942 | SAMN19597777 |
| *spr* | STF | STF_4517_RS | RS | 3/8/2013 | 72 | 3.8 | 7 | 27085325 | 25022314 | SAMN19597778 |
| *pah* | PAHARI/EiJ | PAH.New.1M_LZ | LZ | 7/10/2014 | 75 | 9.5 | 8.2 | 8355637 | 7272288 | SAMN19597779 |
| *pah* | PAHARI/EiJ | PAH.New.1M_RS | RS | 7/10/2014 | 75 | 4.6 | 6.9 | 30810159 | 28255790 | SAMN19597780 |

| | | | | | | | | | | |
|---|---|---|---|---|---|---|---|---|---|---|
| *pah* | PAHARI/EiJ | PAH.New.2M_LZ | LZ | 7/17/2014 | 82 | 11.7 | 8.6 | 23718990 | 20946764 | SAMN19597781 |
| *pah* | PAHARI/EiJ | PAH.New.2M_RS | RS | 7/17/2014 | 82 | 7.3 | 7.5 | 10190552 | 9329587 | SAMN19597782 |
| *pah* | PAHARI/EiJ | PAH.New.4M_LZ | LZ | 7/16/2014 | 81 | 6.4 | 8.3 | 96163007 | 84308080 | SAMN19597783 |
| *pah* | PAHARI/EiJ | PAH.New.4M_RS | RS | 7/16/2014 | 81 | 4 | 7.5 | 10813786 | 9845873 | SAMN19597784 |

**Table S9.** List of genes included in our analyses and whether they were considered expressed, induced, or active in each cell type. Available as a separate supplemental Excel file.

**Table S10.** Number of genes assigned to each regulatory category using both the binomial test and negative binomial test approaches for the fertile F1 mice. Gray boxes indicate the number of genes assigned to the same category using either approach.

| | | **Negative Binomial Test for DE/ASE** | | | | | | | | |
|---|---|---|---|---|---|---|---|---|---|---|
| | | NA | *cis* | *trans* | *cis* X *trans* | compensatory | *cis* + *trans* (opp) | *cis* + *trans* (same) | other | conserved |
| **Binomial Test for DE/ASE** | NA | 0 | 116 | 53 | 0 | 38 | 48 | 26 | 19 | 655 |
| | *cis* | 203 | 200 | 0 | 0 | 0 | 0 | 0 | 43 | 1562 |
| | *trans* | 165 | 0 | 106 | 0 | 0 | 0 | 0 | 0 | 1298 |
| | *cis* X *trans* | 59 | 0 | 38 | 1 | 8 | 0 | 0 | 0 | 860 |
| | compensatory | 67 | 0 | 0 | 0 | 30 | 0 | 0 | 0 | 690 |
| | *cis* + *trans* (opp) | 51 | 0 | 0 | 0 | 149 | 118 | 0 | 0 | 527 |
| | *cis* + *trans* (same) | 94 | 0 | 310 | 0 | 0 | 0 | 127 | 0 | 1029 |
| | other | 250 | 0 | 0 | 0 | 0 | 0 | 0 | 0 | 1493 |

**Table S11.** Read counts from the lapels-suspenders pipeline output for whole testes data.

| Sample | Mack et al. 2016 Sample ID | Total # Reads | Total # Assigned to Parent | LEWES | PWK |
|---|---|---|---|---|---|
| LLWW_SRR2060837 | 148 | 64851628 | 23459524 | 20941397 | 2518127 |
| LLWW_SRR2060842 | 149 | 96823643 | 37201091 | 33111153 | 4089938 |
| LLWW_SRR2060843 | 150 | 85112678 | 32593290 | 29021132 | 3572158 |
| PPCC_SRR2060844 | 151 | 47030803 | 16925427 | 1776081 | 15149346 |
| PPCC_SRR2060846 | 152 | 88496639 | 33850882 | 3394900 | 30455982 |
| PPCC_SRR2060939 | 170 | 68604125 | 26294240 | 2627391 | 23666849 |
| PPLL_SRR2060951 | 52 | 35628710 | 13361236 | 6461350 | 6899886 |
| PPLL_SRR2060955 | 278 | 92806864 | 36956675 | 23978091 | 12978584 |
| PPLL_SRR2060954 | 131 | 23146016 | 8661071 | 4279501 | 4381570 |
| LLPP_SRR2060950 | 93 | 40057612 | 14199347 | 7360932 | 6838415 |
| LLPP_SRR2060952 | 290 | 32200485 | 12058341 | 6153120 | 5905221 |
| LLPP_SRR2060953 | 272 | 112352140 | 43526628 | 22241252 | 21285376 |
